# Neurog1 and Olig2 integrate patterning and neurogenesis signals in development of zebrafish dopaminergic and glutamatergic dual transmitter neurons

**DOI:** 10.1101/2023.05.16.540944

**Authors:** Christian Altbürger, Meta Rath, Wolfgang Driever

**Author notes:** Correspondence should be addressed to: Wolfgang Driever, Institute of Biology 1, Albert-Ludwigs-University Freiburg, Hauptstrasse 1, D-79104 Freiburg, Germany. contributed equally (Christian Altbürger, Meta Rath).

## Abstract

Dopaminergic neurons develop in distinct neural domains by integrating local patterning and neurogenesis signals. While the proneural proteins Neurog1 and Olig2 have been previously linked to development of dopaminergic neurons, their dependence on local prepatterning and specific contributions to dopaminergic neurogenesis are not well understood. Here, we show that both transcription factors are differentially required for the development of defined dopaminergic glutamatergic subpopulations in the zebrafish posterior tuberculum, which are homologous to A11 dopaminergic neurons in mammals. Both Olig2 and Neurog1 are expressed in *otpa* expressing progenitor cells and appear to act upstream of Otpa during dopaminergic neurogenesis. Our epistasis analysis confirmed that Neurog1 acts downstream of Notch signaling, while Olig2 acts downstream of Shh, but upstream and/or in parallel to Notch signaling. Furthermore, we identified Olig2 to be an upstream regulator of *neurog1* in dopaminergic neurogenesis. This regulation occurs through Olig2-dependent repression of the proneural repressor and Notch target gene *her2*. Our study reveals how Neurog1 and Olig2 integrate local patterning signals, including Shh, with Notch neurogenic selection signaling, to specify the progenitor population and initiate neurogenesis and differentiation of A11-type dopaminergic neurons.

## Introduction

The formation of new neurons from neural stem and progenitor cells is a tightly controlled process. During neurogenesis, patterning mechanisms are considered to establish regional transcriptional codes (Vieira et al., 2010), followed by proneural and neural bHLH factors crucial for the progression of neurogenesis (Bertrand et al., 2002). During neurogenesis progression, proneural factors mediate cell cycle exit of progenitor cells and activate downstream targets for terminal differentiation into mature neurons. In this process, proneural genes also contribute to neurotransmitter subtype selection, as for example NEUROG1/NEUROG2 mediate the differentiation of glutamatergic neurons in the dorsal telencephalon and inhibit the activity of the proneural factor ASCL1, which is important for GABAergic neuron formation in the ventral telencephalon (Fode et al., 2000; Schuurmans et al., 2004). Additionally, the neural bHLH factor OLIG2 instructs dorsoventral identity in the spinal cord, as progenitors expressing OLIG2 exclusively give rise to ventrally located motorneurons and oligodendrocytes later on, but not to more dorsally located spinal interneurons (Mizuguchi et al., 2001; Novitch et al., 2001; Zhou and Anderson, 2002).

Understanding dopaminergic (DA) systems development is of great interest, as loss or malfunction especially of DA neurons is associated with severe diseases (Klein et al., 2019). Loss or reduction of DA neurons in the midbrain (mDA) is a hallmark of Parkinson’s disease, while malfunction of A11-type DA neurons has been associated with restless legs syndrome (RLS) and migraine (Clemens et al., 2006). Transcriptional control and neurogenesis of mDA neurons have been extensively studied (Arenas et al., 2015). The proneural factors NEUROG1, NEUROG2 and ASCL1 are expressed in progenitor cells in the ventral midbrain which give rise to mDA neurons (Kele et al., 2006). However, only NEUROG2 is required for the differentiation of mDA neurons, as mice lacking NEUROG2 function show a substantial reduction in mature mDA neurons, whereas earlier precursor cells are unaffected (Andersson et al., 2006; Kele et al., 2006). In vitro studies have shown that NEUROG2 instructs mDA neuron identity in synergy with NR4A2 (Andersson et al., 2007). ASCL1 can partially rescue the loss of mDA neurons in *Neurog2* mutants, but *Ascl1* mutant mice do not exhibit a reduction in mDA neurons. In contrast, Ascl1 is a potent factor in reprogramming fibroblasts into DA neurons (Caiazzo et al., 2011). The neural bHLH factor OLIG2 so far has not been identified to play a role in mDA neurogenesis in mammals. However, the development and differentiation of other DA populations in the mammalian brain has not been extensively studied.

In zebrafish, the proneural factor Neurog1 was identified to have, for specific forebrain DA clusters, a similar function as NEUROG2 for mDA neurons in mammals. It is required for the formation of hypothalamic/posterior tubercular DA neurons and marks their precursor population (Jeong et al., 2006; Russek-Blum et al., 2008). These DA clusters, termed DAC2, 4, 5 and 6 (Rink and Wullimann, 2002), have in common that they produce glutamate as secondary neurotransmitter (Filippi et al., 2014) and are homologous to the OTP-dependent A11 DA neurons in mammals regarding their molecular identity and projections (Ryu et al., 2007; Tay et al., 2011). In this study, we slightly modify the nomenclature for DA neurons in zebrafish previously introduced (Rink and Wullimann, 2002), by using DAC, DA cluster, instead of DC, diencephalic cluster, as some of these neuronal populations reside in in the posterior tuberculum, but others in the hypothalamus and not in the diencephalon (Schredelseker and Driever, 2020). Olig2 was identified to contribute to the neurogenesis of a subset of the A11-type neurons in zebrafish (Borodovsky et al., 2009). Notch signaling is important for the control of DA progenitor populations as perturbations of Notch signaling result in premature differentiation of precursor cells in zebrafish (Mahler et al., 2010). However, it is still unclear how Notch signaling, Neurog1 and Olig2 combine to control DA development. Here, we performed an extensive genetic loss-of-function and overexpression analysis to place Neurog1 and Olig2 into the signaling and transcriptional network that determines ventral forebrain DA development. Our data suggest that Neurog1 and Olig2 integrate patterning (Shh, Nodal, Wnt) and neurogenesis (Notch) signals to control development of all Otp-dependent DA glutamatergic dual transmitter neurons of the posterior tuberculum/hypothalamus, which are homologs of the mammalian A11 group.

## Material and Methods

### Zebrafish maintenance and strains

Zebrafish breeding and maintenance was carried out as described (Westerfield, 1993). Embryos were raised at 28.5°C in egg water containing 0.2 mM 1-phenyl-2-thiourea to prevent pigmentation and staged according to (Kimmel et al., 1995). Larvae were fixed at desired stages with 4% paraformaldehyde. The following published zebrafish lines were used: wildtype AB/TL, *otpa^m866^*; *otpb^sa115^* (Fernandes et al., 2013), *Tg(hsp70:neurog1)ups1* (Madelaine and Blader, 2011), and *neurog1^hi1059^* (Golling et al., 2002).

### Transgenic lines

To generate the *Tg(hsp70l:olig2)* transgenic line, we used gateway cloning followed by transgenesis using the Tol2 transposase system (Kwan et al., 2007). The full-length coding region of *olig2* (ZDB-GENE-030131-4013) was amplified from cDNA by PCR and cloned via BP reaction into the middle donor vector pDONR221 using the following primers: *olig2* forward 5’-GGGGACAAGTTTGTACAAAAAAGCAGGCTCCGCCACCATGGACTCTGACACGAGCC -3’ and *olig2* reverse 5’-GGGGACCACTTTGTACAAGAAAGCTGGGTATTATTTTGAGTCACTGGTCAGCC-3’.

With this *olig2* specific middle entry clone, LR reaction was carried out using the 5’ entry clone p5E-*hsp70l*, the 3’ entry p3E-polyA, and the destination vector pDestTol2CG2 containing the *myl7:eGFP* heart marker. Per embryo 20 pg of the final destination vector pDestTol2CG2;*hsp70-olig2*-pA were injected at 1-cell-stage together with 15 pg Tol2 transposase mRNA, transcribed by SP6 from linearized pCS2FA-transposase vector. The *Tg(hsp70l:olig)m1306* allele (Fig. S1D) was established and used for experiments. Heat-shock-driven *olig2* overexpression was validated in each generation by heat-shock of 26 hpf old embryos, fixation after 90 min, and analysis of heat-shock driven *olig2* expression by whole mount *in situ* hybridization.

The *olig2^m1317^* mutant allele was generated using TALENs cloned with the Golden Gate TALEN kit (Cermak et al., 2011). The targeted site begins at bp 324 after transcription start (ZDB-TSCRIPT-090929-14880) and has the following sequence: 5’-CATGCCCTACGCTCACGGGCCGTCTGTGCGCAAGCTCTCTAAAAT-3’. OLIG2-TALEN1 targets the sequence 5’-CATGCCCTACGCTCA-3’ and OLIG2-TALEN2 5’- CAAGCTCTCTAAAAT-3’ with the RWD sequences HD-NI-NG-NN-HD-HD-HD-NG-NI-HD- NN-HD-NG-HD-NI and NI-NG-NG-NG-NG-NI-NN-NI-NN-NI-NN-HD-NG-NG-NN respectively. The spacer sequence 5’-CGGGCCGTCTGTGCG-3’ between the two TALENs contains a HaeIII restriction site to facilitate genotyping. The mutant line was generated by injecting 100 pg mRNA for each TALEN per embryo. Injected embryos were grown to adulthood and crossed with wildtype fish. Ten embryos per clutch were pooled and lysed. PCR was performed on the lysates, followed by HaeIII restriction digest to screen for clutches containing mutant alleles. Adult F1 fish containing mutant alleles were screened individually by PCR followed by HaeIII restriction digest on gDNA obtained from fin-clips. Uncut PCR product bands were sequenced to identify the exact mutations. The *olig2^m1317^* mutant allele (Fig. S1C) has a 13 bp deletion starting at position bp 341 after transcription start that leads to a frame shift and a premature stop codon at bp 355 – bp 357. The resulting protein has a frame shift after Olig2 amino acid 113 (histidine) and is truncated after 5 additional amino acids. The mutant protein thus lacks the carboxyterminal half of the bHLH domain. The *neurog1^m1469^*mutant allele (Fig. S1B) was generated by CRISPR/Cas9 mediated genome editing. Two sgRNAs flanking the sequence coding for the bHLH domain were designed using CRISPRscan (Moreno-Mateos et al., 2015). The sequences of the sgRNAs including the PAM sequence are GGGAGGACGCGGGGGACGCGGGG and GGCGACGAGGATGCCCCCGACGG. The sgRNA DNA template was generated as described in (Gagnon et al., 2014) and transcribed using the MEGAscript T7 kit (Thermo Fisher Scientific) followed by sodium acetate precipitation. Cas9 mRNA was transcribed from NotI-linearized pCS2+hSpCas9 (Ansai and Kinoshita, 2014) using the mMESSAGE mMACHINE Sp6 kit (Thermo Fisher Scientific). Cas9 mRNA and the two sgRNAs were injected into one cell stage embryos. Injected G0 fish were raised to adulthood and outcrossed with wildtype fish. Five embryos per clutch were pooled and lysed. PCR analysis on the lysates revealed clutches with embryos carrying the desired deletion and were raised to adulthood. Adult F1 fish were screened individually and the PCR products smaller than wildtype were sequenced to identify the mutant alleles carried by the individual fish. The *neurog1^m1469^* allele has lost 396 bp at position bp 96 after translation start site that leads to a protein lacking amino acids 33-164, which contain the whole bHLH domain and surrounding amino acids of the wildtype protein. Amino acids 1-32 and 165-204 are unaffected by the deletion.

### Genotyping

All lines were genotyped by genomic PCR, except *Tg(hsp70l:olig2)m1306* which was identified by *myl7:GFP* expression in the heart. For genotyping the *neurog1^hi1059^* mutant we used published primers (Golling et al., 2002). This genotyping only allows the identification of the mutant allele, as the forward primer is located before and the reverse primer within the viral insertion. To detect the wildtype allele, we used in addition the *neurog1* reverse primer 5’- AGATTGGCCTTTGCTGTCC-3’ located after the viral insertion; thus, two PCRs were carried out per embryo, one to detect the mutant and one for the wildtype allele. The *olig2^m1317^* mutant was genotyped by PCR amplifying a 460bp long piece of DNA flanking the mutated region using *olig2* TALEN forward primer 5’-ATGGACTCTGACACGAGCC-3’ and *olig2* TALEN reverse primer 5’-CTCCGCCGTAGATCTCGCT-3’. The PCR fragment was restricted with HaeIII. Wildtype DNA is cut into two bands while mutant DNA is not cleaved. The three possible genotypes can be clearly identified on a 2.5% agarose gel: wildtype PCR product is cut into 340 bp and 120 bp fragments; heterozygotes show 460 bp, 340 bp and 120 bp fragments, and mutants only 460 bp. *Tg(hsp70:neurog1)ups1* fish were genotyped by genomic PCR using the following primers: *neurog1* forward primer 5’-CCTGACGACACAAAGCTGACC-3’, and the SV40pA reverse primer 5’-CCACTCCCGGACAGACTATCT-3’. The *neurog1^m1469^* mutant was genotyped by genomic PCR using *neurog1* forward primer 5’- TCCTTTTCGCACACGGATGAT-3’ and *neurog1* reverse primer 5’- AGCTGTACACTACGTCGGTT-3’, covering the deleted fragment and amplifying 553 bp in wildtype embryos and 157 bp in homozygous mutant embryos. The *otpa^m866^*, *otpb^sa115^* double mutants were genotyped as described before (Fernandes et al., 2013).

### Heat-shock experiments

Embryos were raised at 28.5°C until they reached the desired stage. For heat-shock, *Tg(hsp70:neurog1)ups1* embryos were incubated for 1 hour at 38.5°C and *Tg(hsp70l:olig2)m1306* embryos for 1 hour at 39°C in a water bath. After heat-shock, the embryos were allowed to develop further at 28.5°C until fixation.

### Whole mount *in situ* hybridization, immunohistochemistry, and TUNEL staining

Whole mount *in situ* hybridization (WISH) (Hauptmann and Gerster, 1994) and fluorescent whole mount *in situ* hybridization (FISH) followed by whole mount immunofluorescence (WIF) were carried out as described before (Filippi et al., 2010). The following probes were synthesized: *her2* (ENSDART00000055709.5)*, her4* (ENSDART00000079274.4; ENSDART00000137573.2; ENSDART00000104209.4; ENSDART00000079265.6)*, her6* (ENSDART00000023613)*, her9* (ENSDART00000078936)*, her12* (ENSDART00000044080), *and her15* (ENSDART00000055707; ENSDART00000055706.6), *neurog1* (Blader et al., 1997), *olig2* (Park et al., 2002), *otpa* (Ryu et al., 2007), *sim1a* (Löhr et al., 2009), *th* (Holzschuh et al., 2001). The templates for the probes for *her2, her4, her6, her9, her12,* and *her15* genes were amplified from genomic DNA by PCR and cloned into the pCRII™-TOPO™ vector using the TOPO™ TA Cloning™ Kit (ThermoFischer Scientific) and sequenced to verify the amplified sequence. TUNEL assay was performed using the ApopTag *in situ* apoptosis detection kit (Chemicon) (Ryu et al., 2005).

### Pharmacological inhibition of signaling pathways

To block Notch signaling, embryos were treated with the γ-secretase inhibitor DAPT (N-[N-(3,5- difluorophenacetyl)-L-alanyl]-S-phenylglycine-t-butylester; Sigma-Aldrich D5942). Treatment was carried out as described before with a final DAPT concentration of 100 µM (Mahler et al., 2010). The final DAPT solution was diluted in 1x E3 medium (5 mM NaCl, 0.17 mM KCl, 0.33 mM CaCl_2_, 0.33 mM MgSO_4_) from a 10 mM stock solution solved in 100% DMSO. Embryos were incubated in DAPT at 24 hpf for 24 hours and fixed at 72 hpf. Control embryos were incubated in 1% DMSO under the same conditions. To inhibit Shh signaling, embryos were treated with Cyclopamine (Merck, CAT-No. 239803) from 24-48 hpf at a final concentration of 50 µM diluted in 1x E3 medium from a 10 mM stock solution dissolved in 100% ethanol. Additionally, ethanol was added to a final concentration of 0.7% to ensure that Cyclopamine stays in solution. Control embryos were incubated in 0.7% ethanol under the same conditions.

### Microscopy, cell counts and statistical analysis

WISH stained embryos were documented using differential interference contrast (DIC) with a Zeiss Axioskop microscope and 10x NA 0.3 or 20x NA 0.5 lenses. Fluorescently labeled embryos stained by WISH and WIF were documented using a Zeiss LSM510 or Zeiss LSM510 META NLO with either a Plan-Apochromat 25x/0.8 Imm Corr or a C-Apochromat 40x/1.1 W Corr objective. For cell counts, embryos were recorded with a Zeiss Axio Examiner D1 using DIC optics and a 20x NA 1.0 lens. Cells were counted in ZEN black 2010 software. Cell numbers of different genotypes were compared using the Wilcoxon-Mann-Whitney rank-sum non-parametric test in GraphPad Prism. Graphs were assembled using Microsoft Excel. Figures were composed with Adobe Photoshop. Levels were linearly adjusted to fill the 8 bit image depth using Adobe Photoshop.

## Results

### *neurog1* and *olig2* mutants reveal DA subtype specific roles

While *neurog1* and *olig2* each have been previously invoked in zebrafish DA differentiation (Borodovsky et al., 2009; Jeong et al., 2006), their specific contributions to early versus late phases of neurogenesis and to distinct anatomical DA groups are not well understood. To dissect the roles of Neurog1 and Olig2 in DA neurogenesis, we analyzed *th* expression in single and double mutant larvae by WISH. While in wildtype at 30 hpf *th* expressing cells of the posterior tuberculum and hypothalamus area are present, in *neurog1^hi1059/hi1059^* homozygous mutants (Fig. S1A) in most embryos no DA neurons are detected at this stage (Fig. 1A and B). At 72 hpf DAC2, DAC4, and DAC5 clusters were largely eliminated, while DAC6 was significantly reduced in neuron number (Fig. 1C – F and M). Thus, Neurog1 appears to be essential for formation of most Otp-dependent DA neurons. All other catecholaminergic (CA) cell clusters had normal cell numbers as judged from *th* expression (Fig. 1M). As the *neurog1^hi1059^* allele is caused by a viral insertion into the 3’ untranslated region of the gene (Golling et al., 2002), the open reading frame is not compromised, and thus, the allele may be a hypomorph. Therefore, using CRISPR/Cas9 mediated genome editing, we generated a new mutant allele, *neurog1^m1469^*, which lacks the coding sequence for the bHLH domain (Fig. S1B). We did not find a significant difference in *th* expression in *neurog1^hi1059^*and *neurog1^m1469^* mutant embryos at 72 hpf (Fig. S2), demonstrating that the mutants represent *neurog1* null alleles. We also generated the *olig2* mutant allele *m1317* via TALEN targeted mutagenesis. *olig2^m1317^* has a 13 bp deletion in the bHLH domain, causing to a frame shift and a preliminary stop codon, and resulting in a truncated version of the Olig2 protein lacking most of the bHLH domain (Fig. S1C). In 30 hpf *olig2^m1317^* homozygous mutant embryos, the number of *th* expressing cells was reduced compared to wildtype siblings (Fig. 1G, H). At 72 hpf, there were significantly less DAC2 and DAC6 neurons, while the number of cells in DAC4 and DAC5, as well as in all other CA clusters appeared normal (Fig. 1I – L and N). Our finding differs from published *olig2* morpholino data (Borodovsky et al., 2009), which reported DAC4 cells to be reduced. Our results indicate that both Neurog1 and Olig2 are essential for proper development of specific DA groups in the hypothalamus and posterior tuberculum.

**Figure 1.**
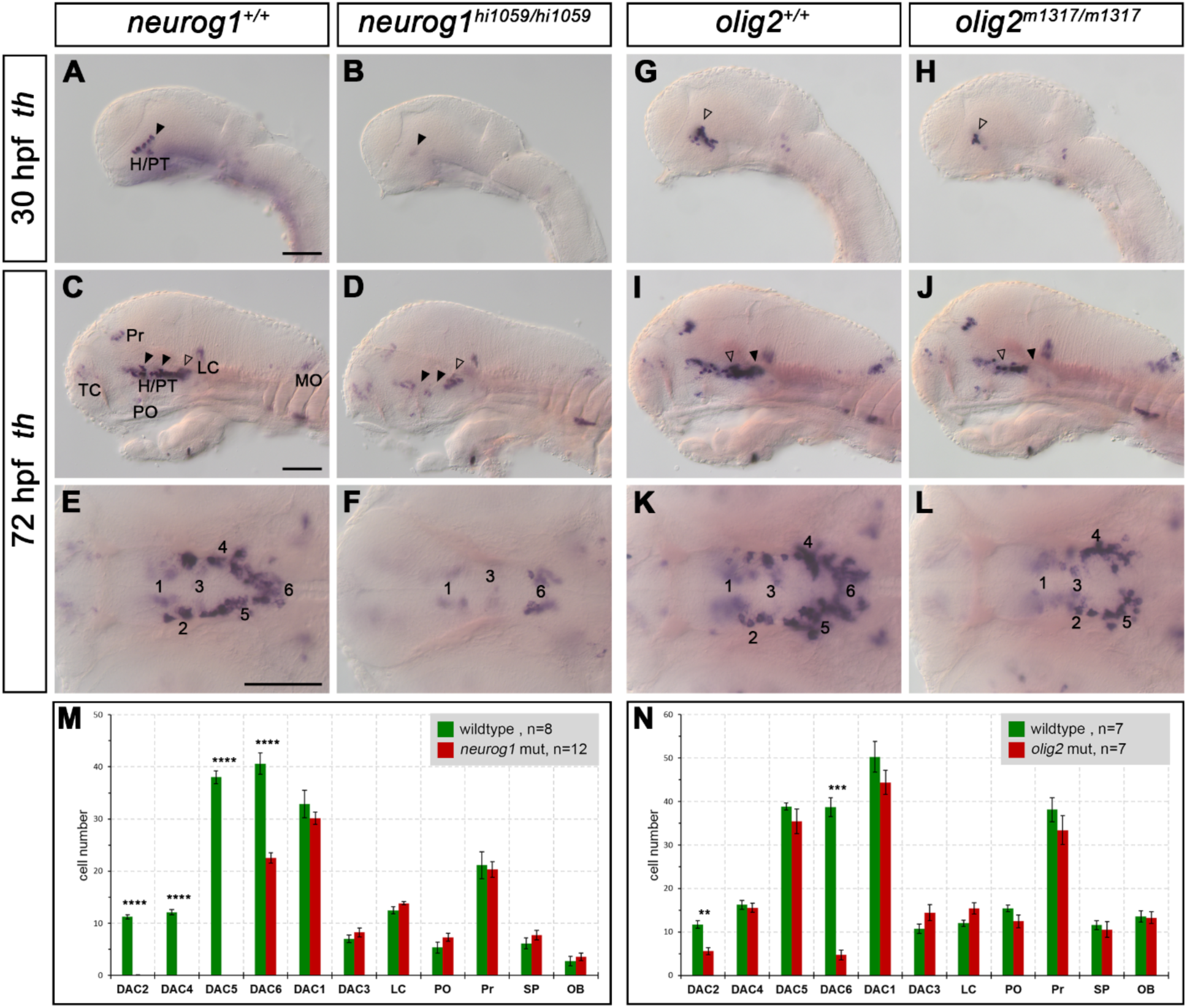
Effect of *neurog1^hi1059/hi1059^*and *olig2^m1317/m1317^* mutants on dopaminergic development. (A-F) *th* expression in (B,D,F) *neurog1^hi1059/hi1059^* mutants compared to (A,C,E) wildtype siblings at (A,B) 30 hpf and (C-F) 72 hpf. (A,B) At 30 hpf in *neurog1^hi1059/hi1059^*mutants no *th+* cells are present (black arrowheads). (C-F) In the H/PT area, DAC2, DAC4 and DAC5 are missing (black arrowheads) and DAC6 is reduced (non-filled arrowhead). (G-L) *th* expression in (H,J,L) *olig2^m1317/m1317^*mutants compared to (G,I,K) wildtype siblings at (G,H) 30 hpf and (I-L) 72 hpf. At 30 hpf in (H) *olig2^m1317/m1317^* mutants, the number of *th+* cells is reduced compared to (G) wildtype (non-filled arrowhead). At 72 hpf in (J,L) *olig2^m1317/m1317^* mutants the numbers of *th+* DAC2 (non-filled arrowheads) and DAC6 (black arrowheads) cells are reduced compared to (I,K) wildtype siblings. (M,N) Quantification of *th+* cells in mutant *neurog1* (M) and *olig2* (N) embryos compared to wildtype siblings at 72 hpf. The number (n) of embryos analyzed for the cell counts are as indicated in the grey box; p-values were calculated using the Mann-Whitney test. p-values indicate significance if they are < 0.05 (**, 0.001 - 0.01; ***, 0.0001 - 0.001; ****, < 0.0001), if no asterisks are present, effects are not significant. For numbers see Supplemental Table 1. Abbreviations: numbers 1 through 7 for DAC1-7, dopaminergic cell cluster 1-6; H, hypothalamus; LC, locus coeruleus; MO, medulla oblongata; OB, olfactory bulb; PO, preoptic area; Pr, pretectum; PT, posterior tuberculum; SP, subpallium; TC, telencephalic cluster. (A-D, G-J) lateral views (enucleated embryos: C, D and I, J) and (E, F, K, L) dorsal views. All images show single optical planes. Anterior is to the left. Scale bars: 100 µm.

To investigate potential overlaps in Neurog1 and Olig2 activity, we analyzed *neurog1^hi1059/hi1059^*; *olig2^m1317/m1317^*double mutants (Fig. 2A, D and I). The double homozygous *th* expression phenotype (Fig. 2D, I) was very similar to the *neurog1* single mutant (Fig. 2B, I), including loss of DAC2, DAC4 and DAC5. However, DAC6 was more severely affected in the double mutants, and the number of remaining DAC6 cells was significantly smaller than in each single homozygous mutant (Fig. 2A – D and I). To determine whether the double mutants may have increased apoptosis, we performed TUNEL staining at 72 hpf (Fig. S3), and detected no difference between *neurog1*; *olig2* double mutants and wildtype siblings. Therefore, the loss of DA neurons is likely caused by deficient DA neurogenesis rather than progenitor or DA neuron death.

**Figure 2.**
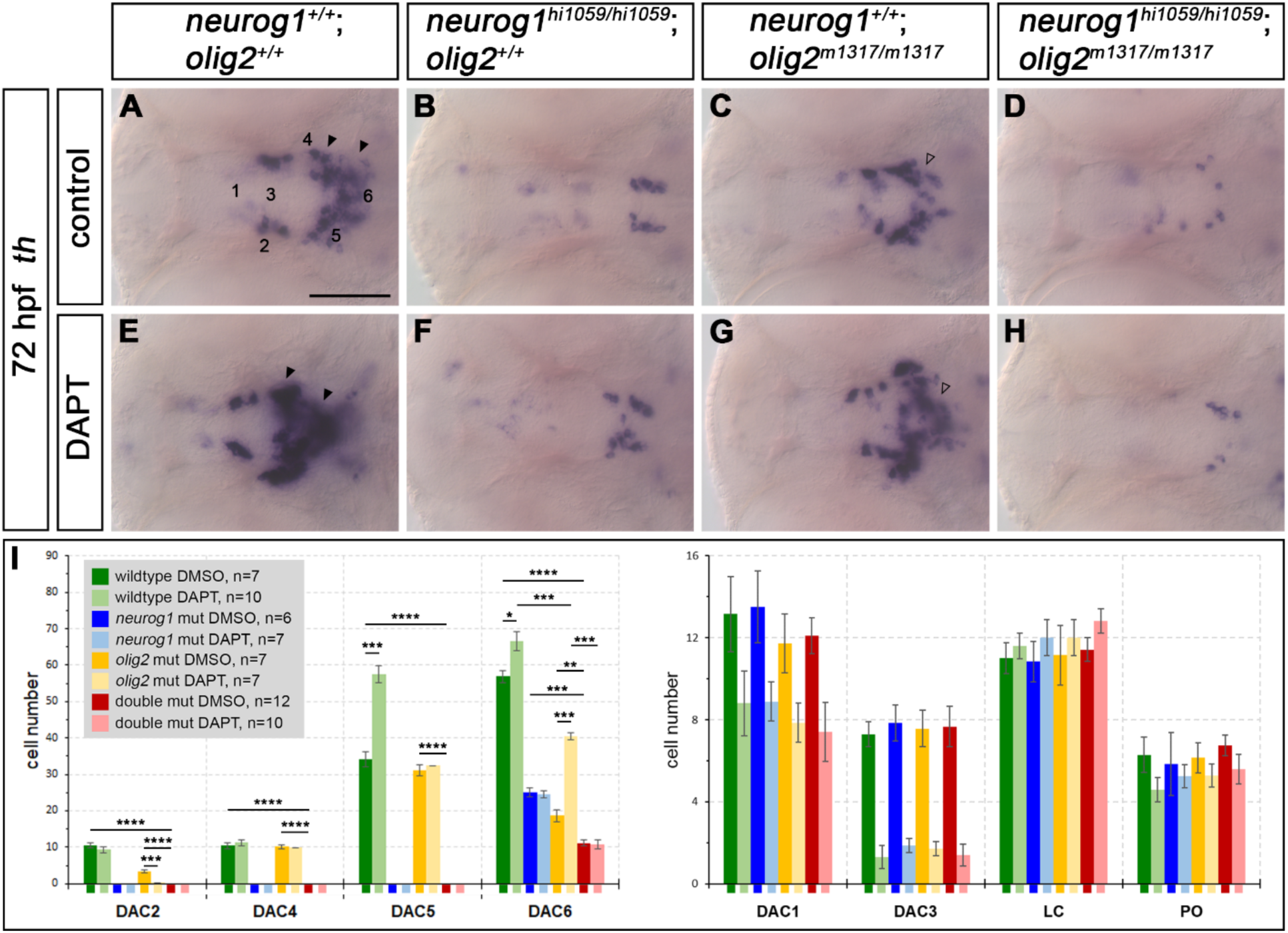
*neurog1* and *olig2* single and double mutant phenotypes and Notch inhibition reveal potential epistatic relationships. (A-H) *th* expression in (A,E) wildtype, (B,F) *neurog1^hi1059/hi1059^* mutants, (C,G) *olig2^m1317/m1317^*mutants and (D,H) *neurog1^hi1059/hi1059^*; *olig2^m1317/m1317^*double mutants in (E-H) DAPT treated embryos compared to (A-D) control siblings at 72 hpf. The numbers 1-6 mark the corresponding (DAC1-DAC6) dopaminergic cell clusters (A). The number of *th+* DAC5 and DAC6 cells is increased in WT embryos (E) after DAPT treatment, compared to (A) control embryos (filled black arrowheads). The number of *th+* DAC6 cells in *olig2^m1317/m1317^* mutant embryos (G) after DAPT treatment is increased compared to control *olig2^m1317/m1317^* mutant embryos (C, open arrowheads). DAPT treatment does not alter the number of *th*+ cells in *neurog1^hi1059/hi1059^*or double mutant embryos (B,F,D,H). (I) Quantification of *th+* cells in single and double mutants compared to wildtype siblings at 72 hpf. The number of embryos analyzed for each genotype are indicated (n-numbers in grey box); p-values were calculated using the Mann-Whitney test. p-values indicate significance if they are < 0.05 (*, 0.01 - 0.05; **, 0.001 - 0.01; ***, 0.0001 - 0.001; ****, < 0.0001). For numbers see Supplemental Table 2. Abbreviations: numbers 1 through 7 for DAC1- 7, dopaminergic cell cluster 1-7; LC, locus coeruleus; PO, preoptic area. All images show single optical planes of dorsal views. Anterior is to the left. Scale bars: 100 µm.

### Olig2 and Neurog1 epistatic position with respect to Notch signaling

To identify the phases during which Neurog1 and Olig2 are active in DA neurogenesis, we treated 24 hpf old embryos with the γ-secretase inhibitor DAPT for 24 hours to inhibit Notch signaling (Geling et al., 2002) (Fig. 2E – H and I). Previous work showed that blocking Notch signaling in wildtype embryos, depending on the time of treatment, leads to an increase or decrease of neurons in specific CA clusters (Mahler et al., 2010). The increase was interpreted as increased neurogenesis caused by suppression of Notch signaling during the lateral inhibition phase of neurogenesis, while the decrease was considered to be a result of depletion of precursor populations when DAPT was applied prior to the main neurogenesis phase of a specific DA group. We chose 24-48 hpf as DAPT treatment window, because DAC1, DAC3, DAC5, and DAC6 neurons undergo phases of active neurogenesis during this period (Mahler et al., 2010). As expected, DAC2 and DAC4 were not affected, as they undergo neurogenesis before the DAPT treatment window. In agreement with Mahler et al., (2010), we found that the clusters DAC5 and DAC6 had increased cell numbers in DAPT treated wildtype embryos compared to control embryos (Fig. 2A, E and I, Fig. 3M). In contrast, DAPT treated *neurog1* mutants did not differ in DAC5 and 6 cell numbers from untreated mutants. Thus, the genetic loss of DA neurons could not be compensated by DAPT treatment (Fig. 2B, F and I, Fig. 3M), consistent with *neurog1* acting downstream of Notch signaling in DA neurogenesis. In *olig2* mutant embryos, DAPT was able to significantly enhance neurogenesis of DAC6 cells (Fig. 2C, G, and I, Fig. 3M), suggesting that Olig2 may act upstream of or in parallel to Notch signaling during development of this DA group. Interestingly, in *olig2* mutants treated with DAPT we did not observe the same increase in DAC5 neurons as in wildtype (Figure 1I), suggesting that Olig2 may also affect development of their progenitors, despite no significant effect of the mutant was observed on DAC5 cell number. The *neurog1*; *olig2* double mutant phenotype could also not be rescued by DAPT treatment (Fig. 2D, H and I, Fig. 3M). These data suggest that nearly all Otp-dependent DA neurons depend on Olig2, Neurog1, or both, except for a small number of caudal DA neurons (potential DAC6 subtype, Fig 2D, H), which in double mutants are also not modulated in number by inhibition of Notch signaling.

**Figure 3.**
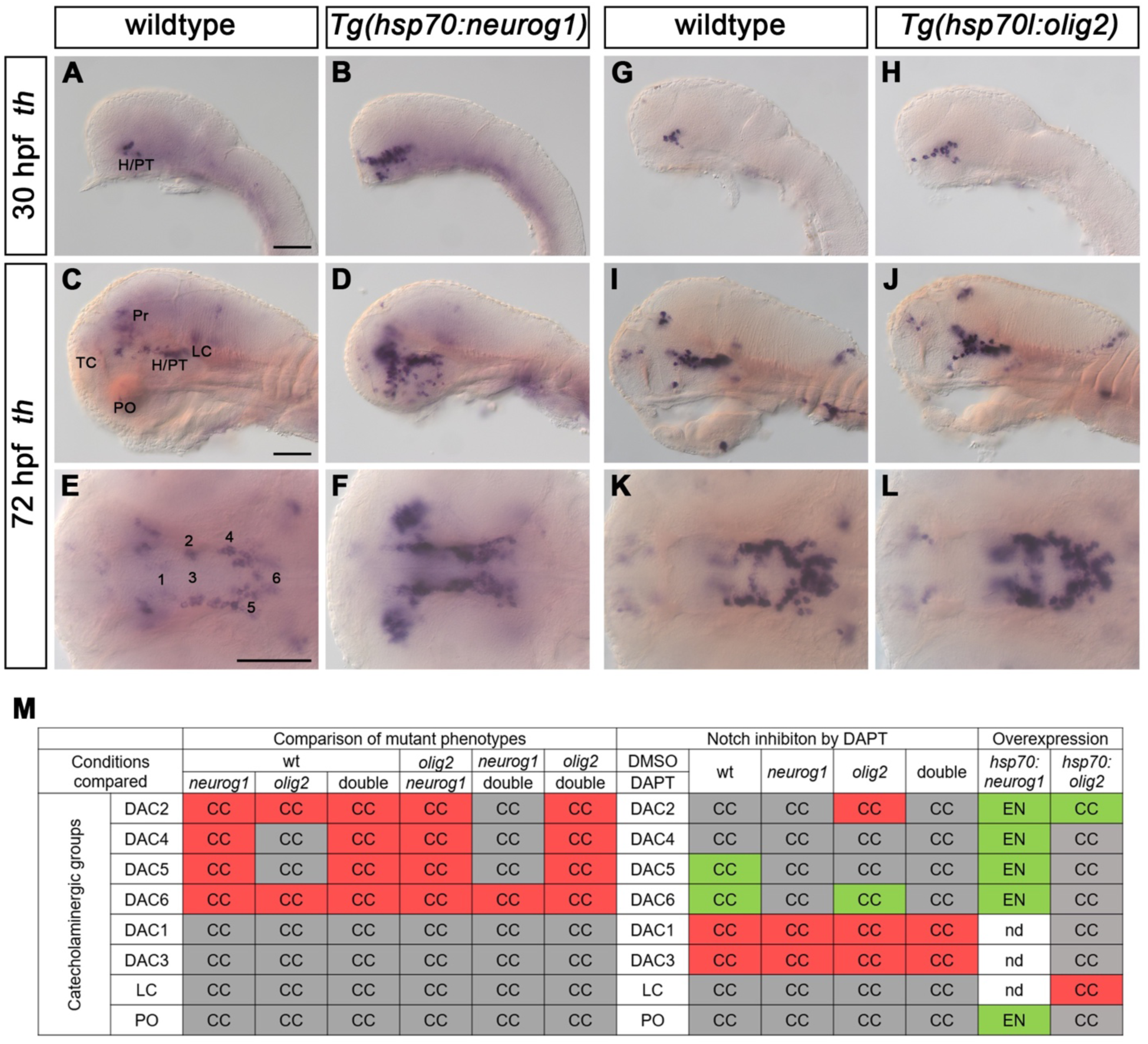
Heat-shock driven overexpression of *neurog1* or *olig2* induces supernumerary and ectopic DA neurons. *th* expression was analyzed following heat-shock in (B,D,F) *Tg(hsp70:neurog1)ups1* compared to (A,C,E) wildtype siblings at (A,B) 30 hpf and (C-F) 72 hpf. (G-L) *th* expression in heat shocked (H,J,L) *Tg(hsp70l:olig2)m1306* compared to (G,I,K) wildtype siblings at (G,H) 30 hpf and (I-L) 72 hpf. The numbers 1 through 6 mark the dopaminergic cell clusters DAC1-DAC6 (E). Overexpression of *neurog1* induces supernumerary and ectopic *th*+ cells at 30 hpf and 72 hpf (B,D,F). Overexpression of *olig2* results in supernumerary *th*+ cells at 30 hpf and 72 hpf, probably belonging to the DAC2 group (H,J,L). (M) Overview of effects of *neurog1* and *olig2* mutant and overexpression, as well as of Notch inhibition on the development of catecholaminergic neurons in the specific anatomical clusters. The specific experiments are shown in Figures 1, 2 and this Figure. CC indicates that cell of the indicated CA groups were counted. EN indicates enhanced stain intensity or ectopic catecholaminergic neurons, which were not counted. Red colored boxes indicate significantly decreased numbers of CA neurons whereas green colored boxes indicate significantly increased cell numbers. Grey colored boxes indicate no change in cell numbers between the given genotypes and treatment conditions. For numbers see Supplemental Table 3. Abbreviations: numbers 1 through 7 for DAC1-7, dopaminergic cell cluster 1-7; H, hypothalamus; LC, locus coeruleus; PO, preoptic area; Pr, pretectum; PT, posterior tuberculum; TC, telencephalic cluster. (A-D, G-J) show lateral views (I,J show enucleated embryos) and (E,F,K,L) dorsal views. All images show single optical planes. Anterior is to the left. Scale bars: 100 µm.

### Overexpression of Neurog1 and Olig2 induces supernumerary DA cells in the diencephalon

To study whether Neurog1 or Olig2 each are sufficient to induce DA cells in vivo, we used transgenic lines expressing *neurog1* or *olig2* under the control of the zebrafish heat-shock promoter *hsp70l* (Fig. S1D). We tested the efficacy of the heat-shock lines by performing a heat-shock at 24 hpf for 45 min at 39°C, followed by WISH for *neurog1* or *olig2* (Fig. S4). For both lines, embryos expressed high levels of transcripts everywhere, and the endogenous expression was not distinguishable from the overexpression anymore, indicating that heat-shock induced levels exceed endogenous expression. Performing heat-shocks at 12 hpf with *Tg(hsp70:neurog1)ups1* embryos resulted in a severe head morphology phenotype, and broader as well as ectopic *th* expression (Fig. 3A – F). Analyzed at 30 and 72 hpf, in hypothalamic, diencephalic and posterior tubercular regions supernumerary cells in DA clusters as well as ectopic *th* expressing cells were found when compared to wildtype siblings (Fig. 3A – F and M). The heat-shock Olig2 overexpression induced *th* expression phenotype was rather mild compared to Neurog1 overexpression. Upon Olig2 heat-shock at 12 hpf, substantially more hypothalamic/posterior tubercular *th* expressing cells could be found at 30 hpf compared to wildtype siblings (Fig. 3G and H). This phenotype was still present at 72 hpf (Fig. 3I – L and M), and may also affect the prethalamic DAC cluster (cells not counted). It is not possible to distinguish non-ambiguously to which DA cluster the additional cells belong. Based on their position, the ectopic cells could be DAC2. Given that this cluster was reduced in *olig2* mutants, we counted them as DAC2 cells (Fig. 3M). Furthermore, the number of DAC6 cells appeared to be unaltered (Fig. 3I – L and M). In summary, Olig2 was sufficient to induce supernumerary DAC2 cells but not DAC6 cells. Additionally, we observed that the noradrenergic neurons of the locus coeruleus are absent in Olig2 overexpressing embryos.

### Distinct and overlapping *neurog1* and *olig2* expression domains

To identify regions in which *neurog1* and *olig2* may be coexpressed or sequentially expressed, we analyzed their expression domains from 18 to 72 hpf by WISH (Fig. S5). The highly dynamic expression patterns suggest potential coexpression in hypothalamus, thalamus, preoptic area and posterior tuberculum. Additionally, we performed double fluorescent *in situ* hybridization (FISH) at 24 and 48 hpf for *neurog1* and *olig2* coexpression analysis (Fig. 4, Supplemental Movies 1, 2). In regions of DA differentiation in the hypothalamus and posterior tuberculum, our FISH confirmed *neurog1* and *olig2* co-expression at a cellular level (Fig. 4). At 30 and 48 hpf, we detected extended *neurog1* and *olig2* co-expression along the ventricular zone (Fig. 4A, C and D, asterisks) as well as at 30 hpf in more lateral regions where Otp-dependent DA precursor are located (Fig. 4A and B, asterisks). A strong overlap of *neurog1* and *olig2* was observed in the hypothalamic/posterior tubercular region where DA neurons develop, and the relative location in the ventricular half of the neuroepithelium suggests that co-expressing cells may be neural progenitors (Supplemental Movies 1, 2). Therefore, we compared the spatiotemporal expression of both genes relative to DA neurons by FISH for *neurog1* or *olig2* expression followed by immunofluorescence for Th as marker for CA neurons (Fig. S6). In the hypothalamus and posterior tuberculum, *neurog1* and *olig2* are expressed in domains medially adjacent to DA neurons that may correspond to precursor territories, but were not co-expressed with Th, and thus not expressed in mature DA neurons (30 hpf: Fig. S6A1 – A3, C1 – C3). This spatial relationship is maintained at 48 hpf, as the expression of both genes is located close to (*neurog1;* Fig. S6B1 – B3) or even directly at the ventricular wall (*olig2;* Fig. S6D1 – D3). In contrast, mature DA neurons marked by Th expression are at more lateral position, strengthening the notion that *neurog1* and *olig2* co-expressing cells may be progenitors.

**Figure 4.**
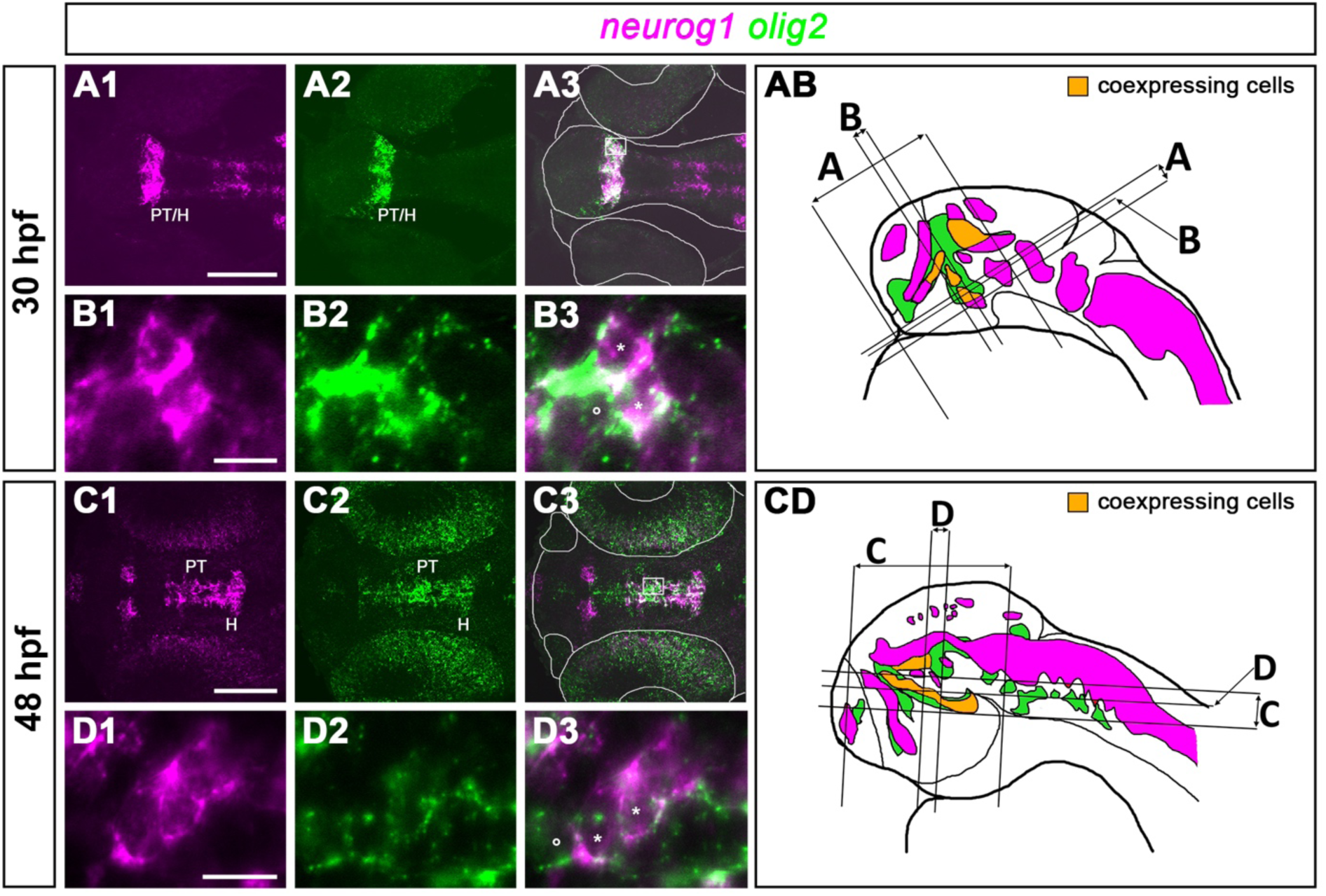
*neurog1* and *olig2* are coexpressed in defined domains during development. (A1-D3) Co-expression of *neurog1* (magenta) and *olig2* (green) at (A1-B3) 30 hpf and (C-D3) 48 hpf. (B1-B3) is a magnification of the corresponding highlighted area of a different embryo in (A3/AB), (D1-D3) of the corresponding highlighted area of a different embryo in (C3/CD). The white line in (A3) frames the outline of the imaged zebrafish larval head, neural tube and eyes; the line in (C3) frames the outline of the imaged zebrafish larval head, nose pits and eyes. At both stages some cells show coexpression of *neurog1* and *olig2* (B3,D3, asterisks). (AB, CD) Schemes show lateral head views of 30 and 48 hpf zebrafish larvae with schematic expression patterns of stained mRNAs and indicate approximate positions of planes and projections. Areas identified to express both genes are colored in orange (see also Supplemental Movies 1, 2). Abbreviations: H, hypothalamus; PT, posterior tuberculum. All images show dorsal views. (B1-B3,D1-D3) show single planes; (A1-A3,C1-C3) are 20-40 µm maximum intensity projections. Anterior is to the left. Scale bars: (A,C) 100 µm, (B,D) 10 µm.

### Olig2 controls subdomains of *neurog1* expression in the hypothalamus

To investigate potential regulatory interactions between Neurog1 and Olig2, we explored *olig2* expression in *neurog1* mutants and *neurog1* expression in *olig2* mutants (Fig. 5). While we did not detect changes in *olig2* expression in the hypothalamus/posterior tuberculum region of *neurog1* mutants at 30 or 48 hpf, we observed a down-regulation of *olig2* in several of its expression domains in the rostral hindbrain at 48 hpf (Fig. 5A – D). In contrast, in *olig2* mutants *neurog1* expression in the hypothalamus/posterior tubercular region was reduced at 30 and 48 hpf, while all main *neurog1* expression domains still formed (Fig. 5E – H). These results indicate that Olig2 is required for proper *neurog1* expression in the hypothalamus/posterior tuberculum, and acts upstream of Neurog1 in this region. To further analyze crossregulation of the two genes, we analyzed *neurog1* expression at 30 hpf in *Tg(hsp70l:olig2)m1306* embryos after heat-shock (Fig. 5I – L). We found that Olig2 overexpression caused an increase in *neurog1* expression in the same expression domain that showed a reduction in *olig2* mutants. Thus, Olig2 is not only required for proper *neurog1* expression in a specific hypothalamic/posterior tubercular region, but also is sufficient to expand *neurog1* expression in vivo in the same region.

**Figure 5.**
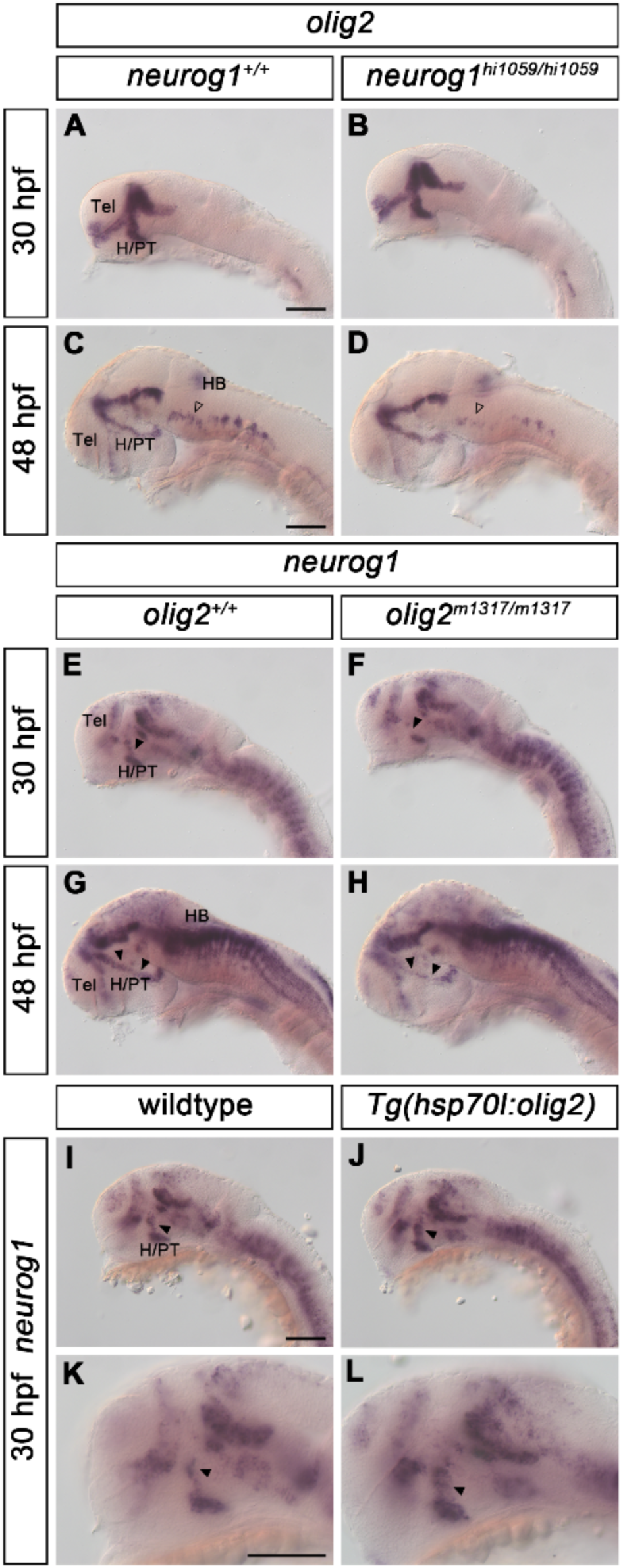
Analysis of potential cross-regulation of *neurog1* and *olig2* expression. (A-D) *olig2* expression in (B,D) *neurog1^hi1059/hi1059^* mutants compared to (A,C) wildtype siblings at (A,B) 30 hpf and (C,D) 48 hpf. At 48 hpf, *olig2* hindbrain expression is reduced in (D) *neurog1^hi1059/hi1059^* mutants compared to (C) wildtype siblings (transparent arrowheads). (E-H) *neurog1* expression in (F,H) *olig2^m1317/m1317^* mutants compared to (E,G) wildtype siblings at (E,F) 30 hpf and (G,H) 48 hpf. At 30 and 48 hpf, *neurog1* expression in the H/PT is reduced in (F,H) *olig2^m1317/m1317^* mutants compared to (E,G) wildtype siblings (black arrowheads). (I-L) *neurog1* expression upon heat-shock in (J,L) *Tg(hsp70l:olig2)m1306* transgenic fish compared to (I,K) wildtype siblings. (K,L) are magnifications of (I,J) respectively. (J,L) *Tg(hsp70l:olig2)m1306* transgenic fish show increased *neurog1* expression in the H/PT when compared to (I,J) wildtype siblings (arrowheads). All *olig2* overexpressing embryos of each genotype showed increased *neurog1* expression in hypothalamus and posterior tuberculum. The phenotypes were observed in N/N (number of embryos with phenotype shown in figure panel / total number of embryos analyzed): (A) 8/8; (B) 6/6; (C) 10/10; (D) 7/7; (E) 7/7: (F) 7/7; (G) 10/10; (H) 8/8; (I,K) 8/8; (J,L) 8/8. Abbreviations: H, hypothalamus; HB, hindbrain; PT, posterior tuberculum; Tel, telencephalon. All images show lateral views (C,D and G,H show enucleated embryos) and single optical planes. Anterior is to the left. Scale bars: 100 µm.

### Olig2 acts upstream of *her2* and downstream of Shh signaling in the posterior tuberculum/hypothalamus

Olig2 has been mainly characterized as a transcriptional repressor (Novitch et al., 2001), suggesting that Olig2 regulation of *neurog1* expression is likely indirect. In the mammalian spinal cord, the regulation of *Neurog2* expression by OLIG2 is mediated via direct repression of *Hes/Her* gene family member expression by OLIG2 (Sagner et al., 2018). We hypothesize a similar mechanism acting in DA neurogenesis in zebrafish. Therefore, we analyzed the expression of the *Hes1/Hes5* zebrafish homologues (Chapouton et al., 2011) in *olig2* mutants. We found the expression of the *Hes1* homologues *her6* and *her9* unchanged in *olig2* mutants compared to wildtype siblings (Fig. S7A - D). For the *Hes5* homologues, we observed no change in expression of *her4*, *her12* and *her15* in *olig2* mutants (Fig. S7E - J). However, the expression of the *Hes5* homologue *her2* is increased in the hypothalamus/posterior tuberculum in *olig2* mutants at 48 hpf (Fig. 6A and B, arrowheads). To further validate this observation and compare the expression of *neurog1* and *her2*, we performed double FISH at 48 hpf on *olig2^m1317/m1317^* mutant embryos. The increase of *her2* expression coincides with a decrease of *neurog1* expression in *olig2* mutants compared to wildtype siblings (Fig. 6C - D’’, white arrowheads). Thus, Olig2 is negatively regulating the expression of *her2* in the hypothalamus/posterior tuberculum, which is positively regulated by Notch signaling (Cheng et al., 2015; Sigloch et al., 2023). To determine whether Shh signaling is involved in regulation of *olig2* expression in the posterior tuberculum/hypothalamus, we inhibited Shh signaling using Cyclopamine from 24 to 48 hpf (Fig. 6E - H). Embryos treated with 50 µM Cyclopamine (Fig. 6F and H) show reduced expression of *olig2* in the posterior tuberculum/hypothalamus when compared to control embryos (Fig. 6E and G). Additionally, the expression of *olig2* in the thalamus (Th) is completely absent in Shh inhibited embryos. Therefore, we conclude that Shh signaling is required for proper regulation of *olig2* expression in the posterior tubercular progenitor domain giving rise to glutamatergic DA neurons.

**Figure 6.**
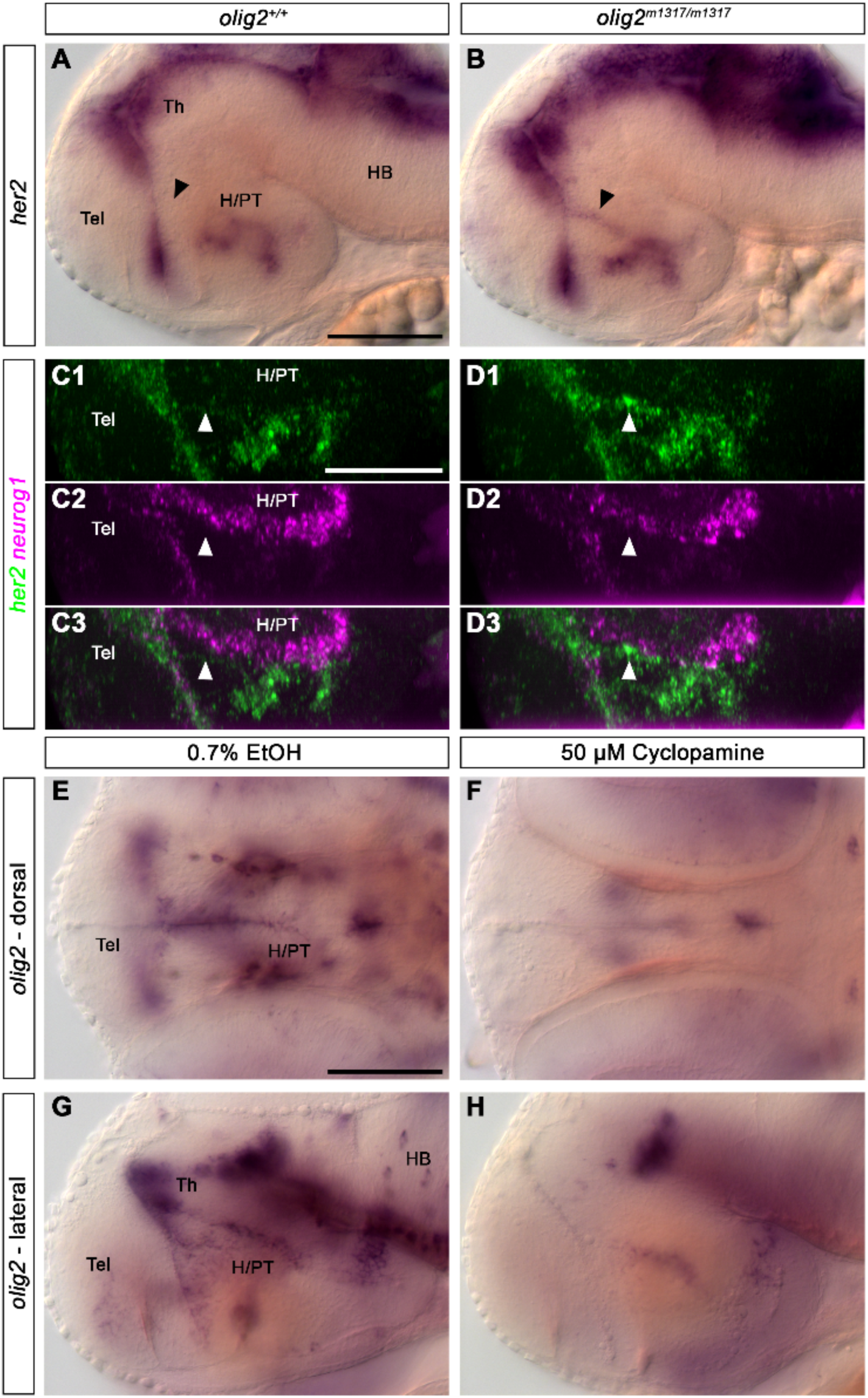
*olig2* expression is regulated by Shh signaling and acts upstream of *her2*. (A, B) *her2* expression in (B) *olig2^m1317/m1317^* mutants compared to (A) wildtype siblings at 48 hpf. The expression of *her2* is increased in the hypothalamus/posterior tuberculum in *olig2^m1317/m1317^* mutant embryos (black arrowheads). (C1-D3) *her2* expression (green) relative to *neurog1* expression (magenta) in (D1-D3) *olig2^m1317/m1317^*mutants compared to (C1-C3) wildtype siblings at 48 hpf. *her2* expression is increased whereas *neurog1* expression is decreased in *olig2^m1317/m1317^*mutant embryos (white arrowheads). (E-H) *olig2* expression at 48 hpf in (F,H) embryos treated with Cyclopamine between 24-48 hpf, compared to (E,G) control embryos treated with ethanol. *olig2* expression is reduced in the hypothalamus/posterior tuberculum and in the thalamus upon inhibition of Shh signaling by Cyclopamine. The phenotypes were observed in N/N (number of embryos with phenotype shown in figure panel / total number of embryos analyzed): (A) 4/4; (B) 7/7; (C1-C3) 4/4; (D1-D3) 4/4; (E,G) 15/15; (F,H) 15/15. Abbreviations: H, hypothalamus; HB, hindbrain; PT, posterior tuberculum; Tel, telencephalon. All images show lateral views, except E, F showing dorsal views; (A and B) show single optical planes; (C1-D3) are 340 µm maximum intensity projections. Anterior is to the left. Scale bars: 100 µm.

### *otpa* expression is reduced in *neurog1* and *olig2* mutants

*otpa* and *otpb* are expressed in progenitors and mature DA neurons in the DAC2, DAC4, DAC5 and DAC6 groups in the hypothalamus/posterior tuberculum, and are strictly required for their development (Ryu et al., 2007). To investigate potential interactions between *neurog1, olig2* and *otpa*, we performed double FISH at 30 hpf (Fig. 7). In the caudal part of the posterior tuberculum, we found co-expression of *otpa* in some cells with *neurog1* or *olig2* (Fig. 7A – E, asterisks). However, the main expression domains of *neurog1* and *olig2* were located closer to the midline, a region where undifferentiated progenitors are located, while *otpa* expression was located more laterally in the mantle zone, where mature DA neurons are located. We next investigated whether *neurog1* or *olig2* might control aspects of *otpa* expression (Fig. 8). At 30 and 48 hpf, *neurog1* mutants showed a reduction in the hypothalamic/posterior tubercular *otpa* expression domains when compared to wildtype siblings (Fig. 8A – D). At 30 hpf, *olig2* mutants also showed reduced *otpa* expression in the hypothalamic/posterior tubercular region compared to controls (Fig. 8E and F), although this effect was weaker when compared to *neurog1* mutants. At 48 hpf, in *olig2* mutants a slight reduction of the caudal part of the hypothalamic/posterior tubercular *otpa* expression domain was detectable (Fig. 8G and H). These results indicate that *neurog1* and *olig2* may indeed act upstream of *otpa*, and that some aspects of the DA cell loss or reduction observed in *neurog1* and *olig2* mutants may be mediated by reduced *otpa* expression level or cell number of *otpa* expressing cells. This may reflect the loss of precursor pools, or, given that *otpa* is continued to be expressed in mature DAC2, 4-6 cluster cells, of mature DA neurons. In addition to *otpa*, we analyzed the expression of the transcription factor *sim1a*, which has previously been shown to be reduced in *olig2* morphants (Borodovsky et al., 2009), and to act in parallel to *otpa* in the terminal differentiation of DA neurons (Löhr et al., 2009) (Fig. 8I - P). In *neurog1* mutants, *sim1a* expression is reduced at 30 hpf in the hypothalamus/posterior tuberculum (Fig. 8I and J, arrowheads), whereas it appears unchanged at 48 hpf (Fig. 8 and L, arrowheads). Similar to *olig2* morphants, *olig2* mutants exhibit a reduction in *sim1a* expression in the hypothalamus/posterior tuberculum at 30 hpf (Fig. 8M and N, arrowheads) as well as at 48 hpf (Fig. 8O and P, arrowheads). To determine if early Otpa/b function may also be involved in the transcriptional regulation of *olig2*, we analyzed *olig2* expression in *otpa^m866^*, *otpb^sa115^*mutant embryos at 48 hpf (Fig. S8). No change in *olig2* expression in the posterior tuberculum/hypothalamus was observed in *otpa*, *otpb* mutants (Fig. S8B) in comparison to wildtype siblings (Fig. S8A). Therefore, we conclude that Olig2 regulates aspects of *otpa* expression in late DA progenitors and DA neurons, but not vice versa.

**Figure 7.**
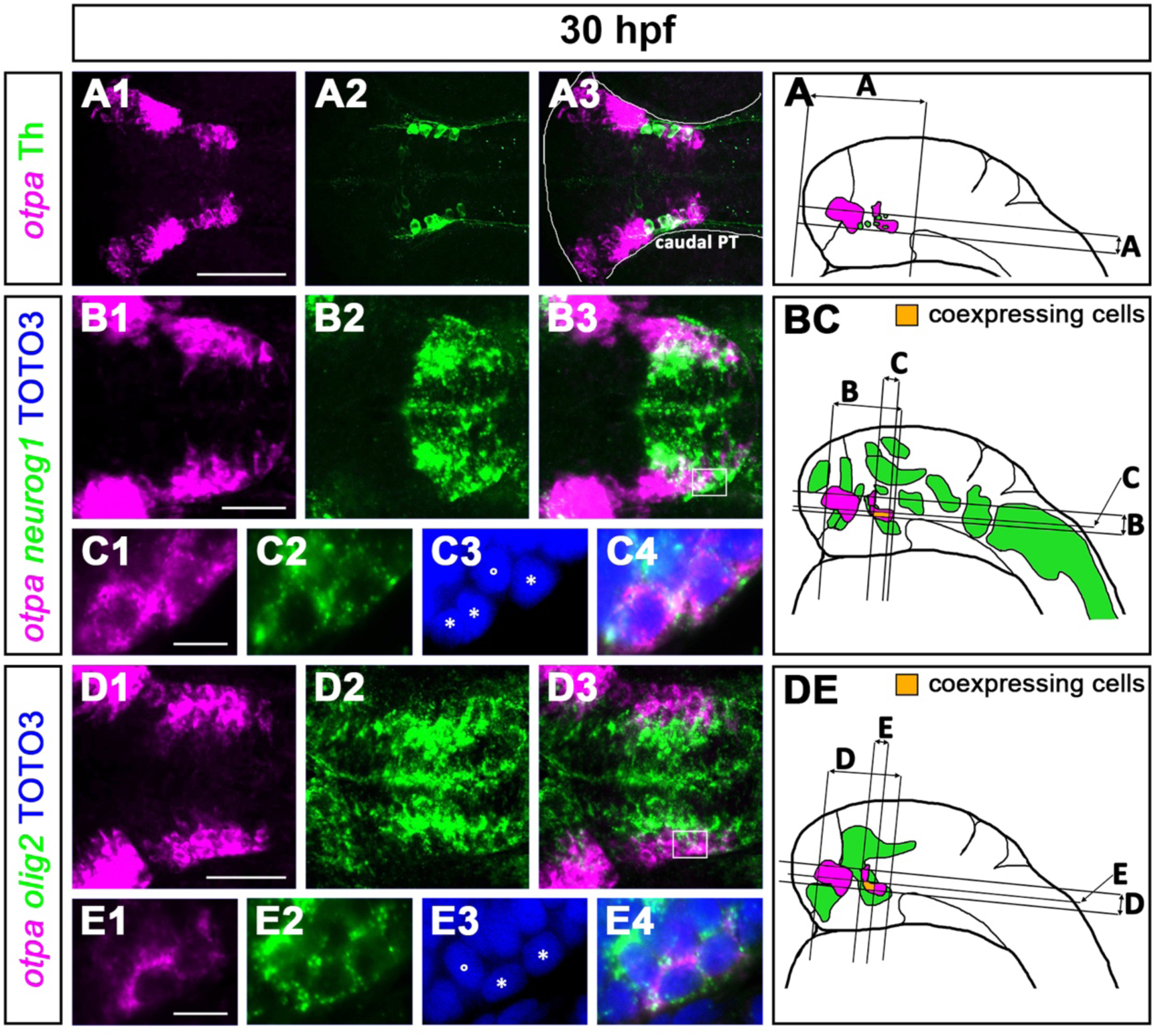
*neurog1* and *olig2* are coexpressed with *otpa.* (A1-A3) *otpa* mRNA expression relative to Th protein expression. (B1-C4) co-expression of *otpa* (magenta) and *neurog1* (green). (D1-E4) co-expression of *otpa* (magenta) and *olig2* (green). (C1-C4) is a magnification of the corresponding highlighted area of a different embryo in (B3/BC), (E1-E4) of the corresponding highlighted area of a different embryo in (D3/DE). The white line in (A3) frames the outline of the imaged zebrafish larval head and eyes. C3,4 and E3,4 show TOTO3 nuclear stain. (A, BC, DE) Schemes show lateral head views of 30 hpf zebrafish larvae with schematic expression patterns of stained mRNAs and proteins and indicate approximate positions of planes and projections. Areas with coexpression are colored in orange. Abbreviations: PT, posterior tuberculum. All images show dorsal views. (C1-C4,E1-E4) show single confocal planes; (A1-B3, D1-D3) are 30-40 µm maximum intensity projections. Anterior is to the left. Scale bars: (A) 100 µm, (B,D) 50 µm, (C,E) 10 µm.

**Figure 8.**
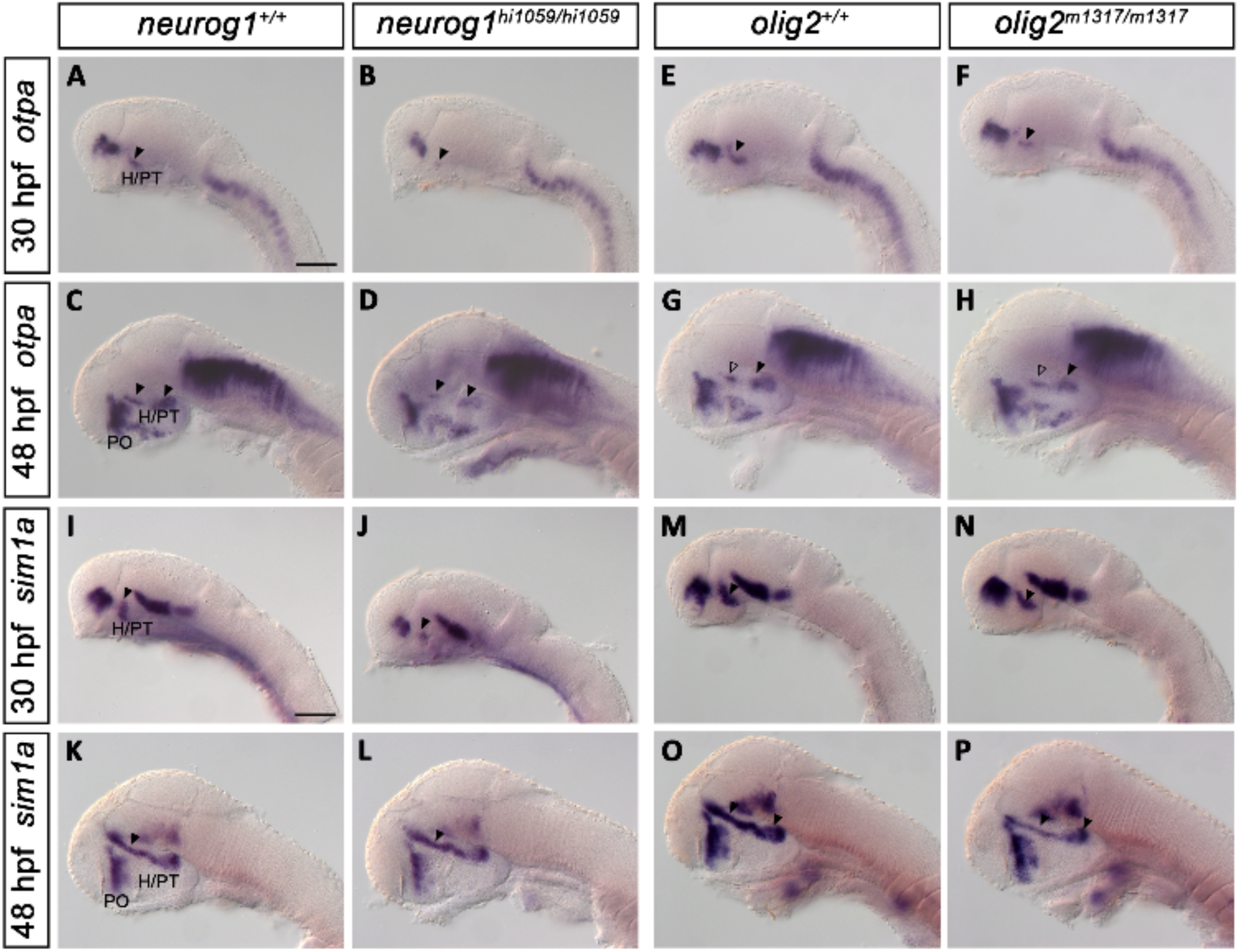
*otpa* and *sim1a* expression is reduced in the H/PT region upon loss of either Neurog1 or Olig2. (A-D) *otpa* expression in (B,D) *neurog1^hi1059/hi1059^* mutants compared to (A,C) wildtype siblings at (A,B) 30 hpf and (C,D) 48 hpf. In *neurog1^hi1059/hi1059^* mutants *otpa* expression in the H/PT is reduced if compared to wildtype siblings (black arrowheads). (E-H) *otpa* expression in (F,H) *olig2^m1317/m1317^*mutants compared to (E,G) wildtype siblings at (E,F) 30 hpf and (G,H) 48 hpf. At 30 hpf in (F) *olig2^m1317/m1317^*mutants, the *otpa* expression in the H/PT is reduced compared to (E) wildtype (black arrowhead). At 48 hpf in (H) *olig2^m1317/m1317^* mutants *otpa* expression in the caudal H/PT is reduced (black arrowheads) and unchanged in the rostral H/PT (non-filled arrowheads). (I-L) *sim1a* expression in (J,L) *neurog1^hi1059/hi1059^* mutants compared to (I,K) wildtype siblings at (I,J) 30 hpf and (K,L) 48 hpf. In *neurog1^hi1059/hi1059^*mutants *sim1a* expression in the H/PT is reduced if compared to wildtype siblings at 30 hpf but not at 48 hpf (black arrowheads). (M-P) *sim1a* expression in (N,P) *olig2^m1317/m1317^* mutants compared to (M,O) wildtype siblings at (M,N) 30 hpf and (O,P) 48 hpf. In *olig2^m1317/m1317^*mutants, the *sim1a* expression in the H/PT is reduced compared to wildtype at 30 hpf as well as 48 hpf (black arrowheads). The phenotypes were observed in N/N (number of embryos with phenotype shown in figure panel / total number of embryos analyzed): (A) 10/10; (B) 15/15; (C) 9/9; (D) 12/12; (E) 18/18: (F) 12/12; (G) 17/17; (H) 6/6; (I) 6/6; (J) 7/7; (K) 5/5; (L) 10/10; (M) 4/4; (N) 11/11; (O) 3/3; (P) 6/6. Abbreviations: H, hypothalamus; PO, preoptic area; PT, posterior tuberculum. All images show lateral views (48 hpf: enucleated embryos) and are single planes. Anterior is to the left. Scale bar: 100 µm.

## Discussion

Patterning as well as neurogenesis signals controlling DA development have been intensively studied for ventral tegmental DA neurons, but how both types of signals are integrated to define position and size of forebrain DA neuron clusters is less well understood. Zebrafish develop DA neurons exclusively in the forebrain, with distinct anatomical DA clusters having specific co-transmitter phenotypes and axons targeting distinct regions of the brain (Filippi et al., 2014; Rink and Wullimann, 2002; Tay et al., 2011). Here, we investigate how the bHLH factors Neurog1 and Olig2 contribute to zebrafish DA neurogenesis. We find that all DA clusters with a glutamatergic cotransmitter phenotype depend on Neurog1 activity, while the DAC6 cluster in addition depends on Olig2 activity. Shh controls Olig2 expression, which in turn represses Her2, a repressor of neurogenin, to control DAC6 development. Thus, Olig2 relays Shh patterning information into control of DAC6 neurogenesis. Together, Neurog1 and Olig2 integrate patterning information from Nodal, Wnt and Shh signaling, as well as Notch signaling, to determine position and size of the ventral forebrain glutamatergic DA clusters DAC2, 4, 5 and 6 (Figure 9).

**Figure 9.**
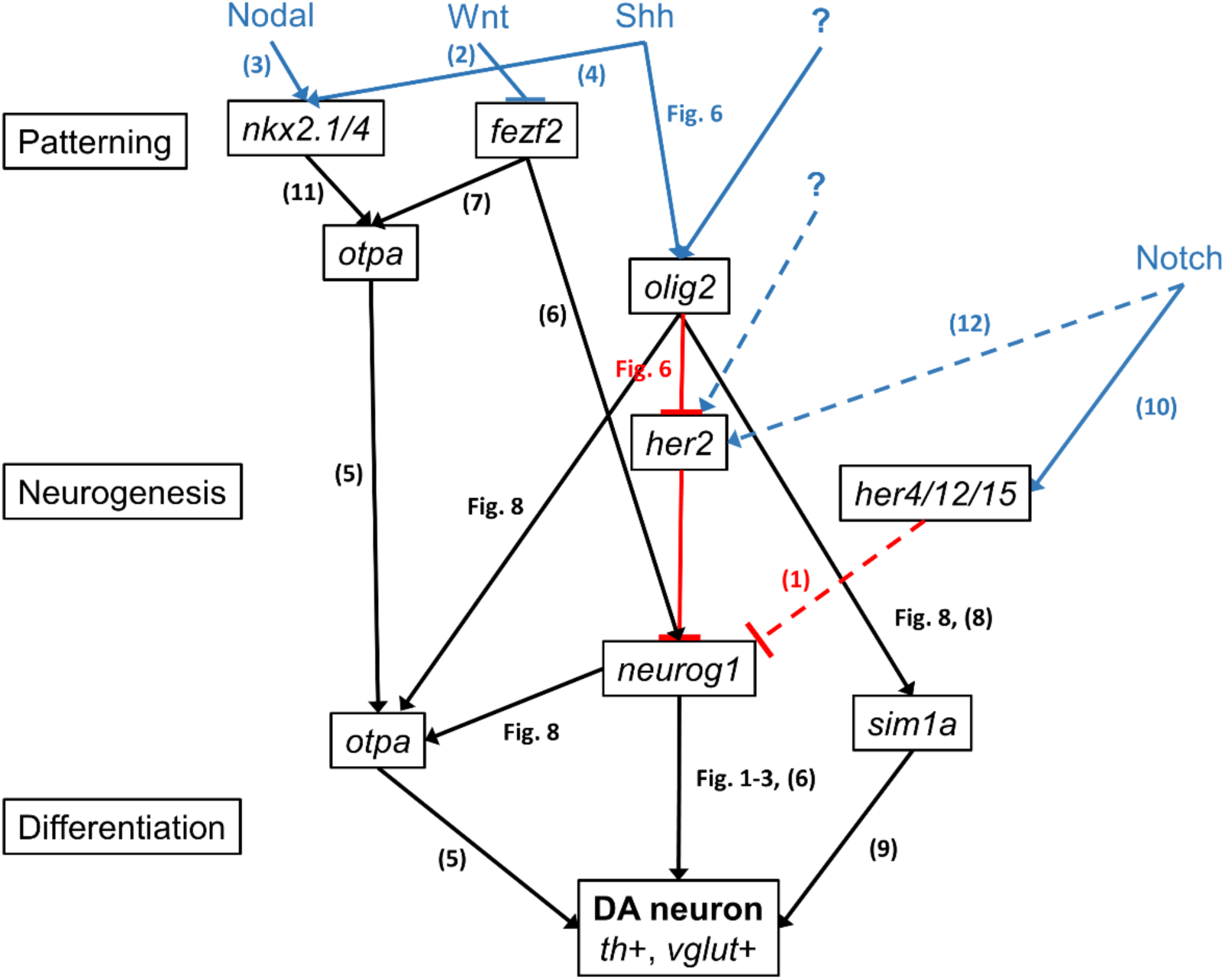
Transcriptional network integrating patterning and neurogenesis signals to control posterior tubercular / hypothalamic A11-type DA neuron development in zebrafish. The network model shows *olig2* and *her2* as components that selectively contribute to DAC6 cluster development, while all other components control DAC2, 4, 5 and 6 cluster development, which are the only glutamatergic DA clusters in zebrafish embryos. Published evidence (numbers in brackets refer to references listed below) reveals Nodal, Wnt and Shh patterning signals to control early *nkx2.1/4, fezf2* and *otpa* expression in progenitors. In DAC6, Olig2 is regulating the expression of *neurog1* through inhibition of *her2*, which is a Notch-dependent *Hes/Her* gene. Olig2 and Neurog1 regulate the expression of *otpa,* which is crucial for the formation of the A11- type DA neurons, whereas *sim1a* is mainly regulated by Olig2 alone. Shh signaling contributes to the control of *olig2* expression. Black arrows indicate positive genetic interactions. Red lines indicate inhibitory genetic interactions. Blue arrows indicate signaling pathways contributing to control of gene expression. Dashed lines indicate partial or unclear genetic/signaling interactions. References: (1) Takke et al., 1999; (2) Hashimoto et al., 2000; (3) Rohr et al., 2001; (4) Tyurina et al., 2005; (5) Ryu et al., 2007; (6) Jeong et al., 2006; (7) Blechman et al., 2007; (8) Borodovsky et al., 2009; (9) Lohr et al., 2009 ; (10) Chapouton et al., 2011; (11) Manoli and Driever, 2014; (12) Cheng et al., 2015.

A role of Neurog1 in the formation of hypothalamic/posterior tubercular DA neurons has previously been described based on morpholino knockdown of *neurog1* (Jeong et al., 2006). We confirmed this finding genetically, both with the *neurog1^hi1059^* retroviral insertion allele that leaves the Neurog1 ORF intact (Golling et al., 2002; Yang et al., 2012), and our new CRISPR/Cas9 induced *neurog1^m1469^* null allele. The role of Neurog1 in the formation of A11-type neurons appears similar to the function of NEUROG2 in mammalian midbrain DA neurogenesis (Andersson et al., 2006; Kele et al., 2006). Here, NEUROG2 is involved in neurogenic selection of mDA neurons from progenitor pools, as well as, together with other transcription factors like NR4A2, in the differentiation of DA neurons (Andersson et al., 2006; Andersson et al., 2007). In contrast, mouse *Neurog1* is expressed in the progenitor zone giving rise to mDA neurons, but is not required for their differentiation (Kele et al., 2006). Zebrafish do not have a *Neurog2* homologue, and therefore Neurog1 is postulated to exhibit NEUROG1 and NEUROG2 function. The finding that *neurog1* overexpression from mRNA injections (Jeong et al., 2006), as well as our heat-shock driven overexpression, induce ectopic DA differentiation, suggests that also in zebrafish, *neurog1* contributes to differentiation of DA neurons. In the development of the mammalian telencephalon NEUROG1 and NEUROG2 are mediating the specification of pallial progenitors into glutamatergic neurons while repressing the formation of GABAergic neurons (Britz et al., 2006; Fode et al., 2000; Schuurmans et al., 2004). Furthermore, *Neurog1*/*2* expression marks thalamic progenitor cells giving rise to glutamatergic neurons (Vue et al., 2007). The fact that the A11-type DA neurons, which we found to be lost or severely reduced in *neurog1* mutants, are glutamatergic (Filippi et al., 2014), supports the hypothesis that the important role of NEUROG1/NEUROG2 in glutamatergic neurogenesis is evolutionary conserved in vertebrates. This is further supported by the observation that *neurog1* morphant zebrafish have reduced glutamatergic neurons in the thalamus (Scholpp et al., 2009).

The neural bHLH factor OLIG2 has not been reported to be involved in the formation of mDA neurons in mammals. OLIG2 function is well characterized for spinal cord motorneurons, and it is crucial for oligodendrocyte formation, both developing from the ventral spinal cord primary motor neuron (pMN) domain (Lu et al., 2002; Meijer et al., 2012; Zhou and Anderson, 2002). Olig2 has also been described to contribute to a subset of A11-type DA neurons in the hypothalamus/posterior tuberculum via promotion of *sim1a* expression in zebrafish (Borodovsky et al., 2009), and to glutamatergic neurons in the hypothalamus in mouse (Ono et al., 2008). In our experiments, global overexpression of Olig2 does induce additional DA neurons only in regions of ongoing DA neurogenesis, revealing its limited potential to drive ectopic DA differentiation. In addition to the reduction of DA neurons in *olig2* mutants, we observe a reduction of *neurog1* expression in the region of the DA precursor population. Therefore, we propose that Olig2 regulates *neurog1* expression directly or indirectly. This idea is supported by our finding that Olig2 overexpression results in increased *neurog1* expression. However, as Olig proteins are mainly acting as repressors (Mizuguchi et al., 2001; Novitch et al., 2001), the regulation of *neurog1* expression is likely indirect. A similar mechanism has been shown for pMN progenitor development in the spinal cord. pMN progenitors express OLIG2, which promotes the expression of *Neurog2*, which in consequence initiates cell cycle exit and differentiation into mature motorneurons (Mizuguchi et al., 2001; Novitch et al., 2001). Similar to pMN progenitors (Mizuguchi et al., 2001), we find *olig2* and *neurog1* coexpressed in progenitor cells that will give rise to A11-type DA neurons.

Notch signaling contributes to A11-type DA neuron formation in zebrafish through maintaining progenitor pools for continuous neurogenesis of DA neurons (Mahler et al., 2010). Mutants for the Notch ligands *dla* and *dld* show that, after enhanced early neurogenesis, loss of Notch signaling as development progresses results in loss of DA precursor cells expressing *neurog1* and *olig2* (Mahler et al., 2010). In our study, we pharmacologically inhibited active Notch signaling through DAPT between 24-48 hpf, and found in wildtype embryos the expected increase in DAC5 and DAC6 neurons resulting from increased neurogenesis. In contrast, in *neurog1* mutants no increase was observed, confirming that Notch signaling acts upstream of *neurog1*, and that Neurog1 is required for DAC5 and DAC6 neurogenesis. In contrast, blocking Notch signaling can partially rescue the *olig2* mutant phenotype, suggesting that Olig2 acts upstream or in parallel to Notch signaling. Recently, it was shown that murine pMN progenitors which have high levels of OLIG2 protein have low levels of the proneural repressor and Notch target genes *Hes1/Hes5* (Sagner et al., 2018). *Olig2* mutants have higher HES5 levels in pMN progenitor cells, whereas *Neurog2* expression is downregulated. *Olig2* expression in turn is induced by Shh signaling emanating from the floor plate of the spinal cord (Lu et al., 2000). Thus, the regulatory network of OLIG2, HES5 and NEUROG2 integrates patterning and neurogenesis signals to control motor neuron pool size. We suggest that a similar mechanism acts in hypothalamic/posterior tubercular DA neurogenesis in zebrafish. When we screened zebrafish *HES/HER* homologues (Chapouton et al., 2011), we found the expression of the *Hes5* homologue *her2* to be controlled by Olig2 in the posterior tubercular DA progenitor region. Interestingly, the expression of *her15*, which is clustered genetically with *her2*, is not affected in *olig2* mutant embryos, suggesting that Olig2 specifically regulates only one *Hes5* homologue in zebrafish. Furthermore, we observed that Shh signaling contributes to control of *olig2* expression in the hypothalamus/posterior tuberculum. As Shh signaling is only partially regulating *olig2* expression in the DA progenitor domain, additional signaling pathways might contribute to its regulation. A good candidate is Nodal signaling which regulates the expression of *nkx2.1* and *nkx2.4a/b* to instruct ventral forebrain patterning (Manoli and Driever, 2014; Rohr et al., 2001). Nodal and Shh signaling both have been previously shown to be required for ventral forebrain DA development (Holzschuh et al., 2003). Thus, Nodal and Shh signaling could act synergistically to regulate *olig2* expression similar to the *nkx2.1/4* genes (Karlstrom et al., 2003; Manoli and Driever, 2014; Tyurina et al., 2005), or *olig2* could also be a downstream target of Nkx2.1/4.

The differentiation of A11-type DA neurons relies on the transcription factor Otp (Del Giacco et al., 2006; Fernandes et al., 2013; Ryu et al., 2007). Fezf2 as well as Nkx2.1/4 regulate the expression of *otpa* and *otpb* in the hypothalamic/posterior tubercular region that gives rise to A11- type DA neurons (Blechman et al., 2007; Manoli and Driever, 2014; Wolf and Ryu, 2013). Furthermore, it was shown that Fezf2 acts upstream of Neurog1 in DA neurogenesis in zebrafish (Jeong et al., 2006). Canonical Wnt signaling represses the expression of *fezf2*, thereby attaining the correct number of DA neurons (Hashimoto et al., 2000; Russek-Blum et al., 2008). Blechman et al. (2007) proposed that Neurog1 might be regulating *otp* expression directly. We found both *neurog1* and *olig2* to be coexpressed with *otpa* in a group of cells in the mantle zone. These cells are most probably late progenitor cells, which start to differentiate into neurons. We observed a specific reduction of *otpa* expression in the hypothalamus/posterior tuberculum in *neurog1* mutants, while *otpa* expression was reduced to a lesser extend in *olig2* mutants. Therefore, both bHLH factors appear to regulate aspects of *otpa* expression. With respect to DA neurogenesis, Otp has two important activities: first it is a read-out of patterning information that defines the DA progenitor territory, and second it drives DA differentiation and continues to be expressed in mature DA neurons, where it also maintains its own expression (Del Giacco et al., 2006; Fernandes et al., 2013; Ryu et al., 2007). The analysis of *otpa^m866^*, *otpb^sa115^* double mutants revealed that Otp activity acts strictly downstream of Olig2, and Otp does not affect *olig2* expression.

In summary, we show that the bHLH factors Neurog1 and Olig2 differentially contribute to hypothalamic/posterior tubercular DA neurogenesis (Figure 9). While Neurog1 is important for the formation of all Otp-dependent A11-type glutamatergic DA neurons, Olig2 is only required for the DAC6 cluster. Olig2 inhibits *her2* expression, and thus indirectly derepresses *neurog1* expression in progenitor cells. Olig2 contributes to Otp and Sim1, and Neurog1 to Otp expression, together specifying the DA phenotype of the neurons, while in parallel Neurog1 may also specify the glutamatergic dual transmitter phenotype. In essence, Olig2 and Neurog1 integrate information provided by patterning signals (Nodal, Shh, Wnt) and neurogenic selection by Notch signaling to specify position and size of the hypothalamic/posterior tubercular DA neuron clusters.

## Contributions

WD, MR and CA designed the study. MR and CA performed experiments and analyzed the data. MR and CA wrote the manuscript and assembled the figures. WD edited the manuscript, obtained funding and supervised the project.

## Acknowledgements

We thank the Life Imaging center for technical support, T. Schredelseker, M. Smirnov, A. M. Fernandes and A. Filippi for critical discussions and S. Götter for excellent zebrafish care. We thank Beatrice Weber and Christian Sigloch for cloning and providing the vectors used to synthesize the WISH probes against the *her* genes. We thank Patrick Blader for giving us the *Tg(hsp70:neurog1)ups1* transgenic line. The *neurog1^hi1059^*mutant allele was obtained from the Zebrafish International Resource Center (ZIRC). This study was supported by the German Research Foundation (DFG) under Germanýs Excellence Strategy (CIBSS - EXC 2189 – Project ID 390939984) and Excellence Initiative (BIOSS - EXC 294) as well as by the EC programs EU- FP7 IP ZF-HEALTH, and FP7 IP DOPAMINET). The authors declare no conflict of interest.

