## Supplemental Materials for "Neurog1 and Olig2 integrate patterning and neurogenesis signals in development of zebrafish dopaminergic and glutamatergic dual transmitter neurons"

Supplemental Figures 1 to 8 (Pages 2-9)

Supplemental Videos 1 to 2 (Legends Page 10)

Supplemental Tables 1 – 3 (Pages 11 – 19)

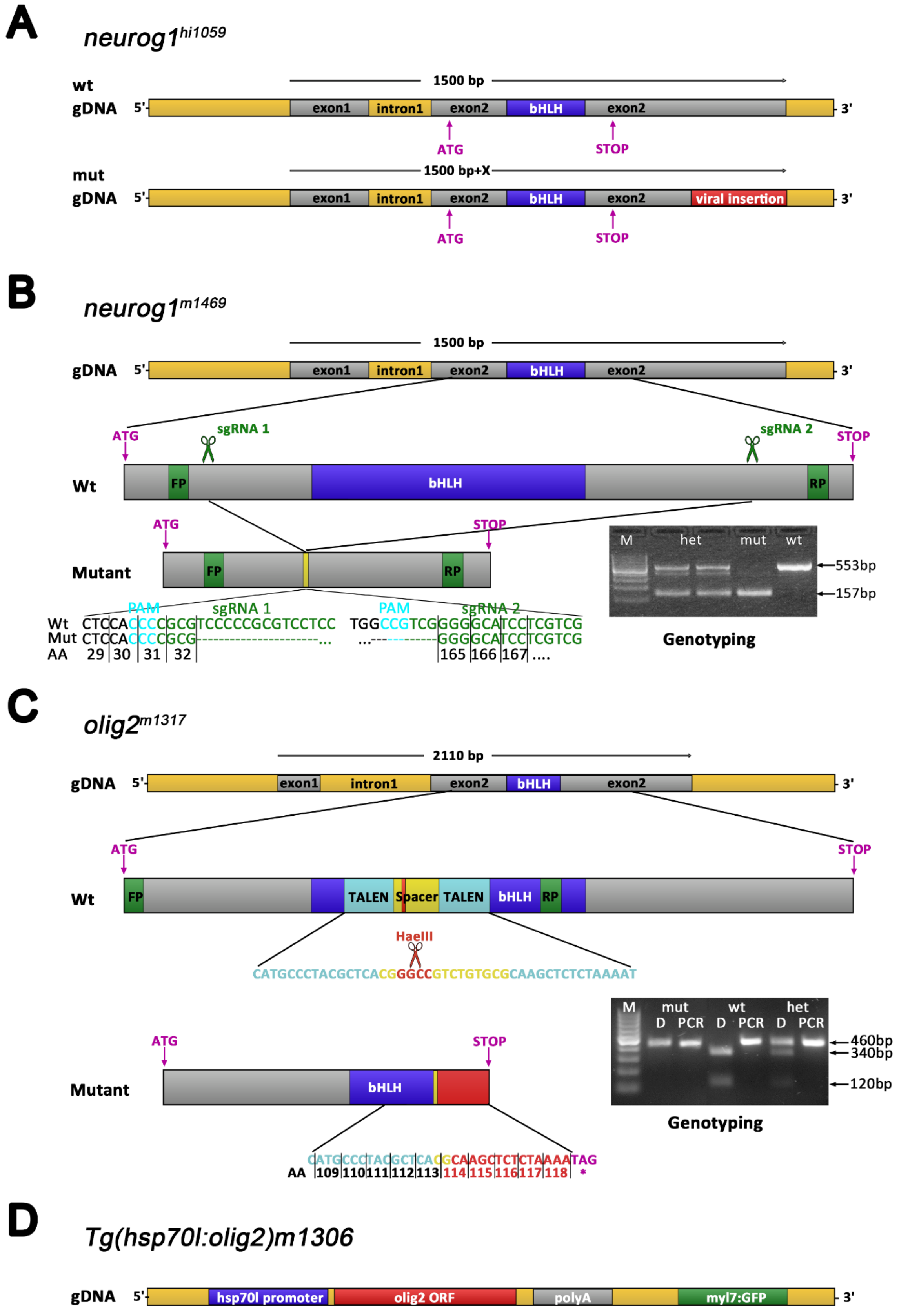


**Figure S1. Overview of genetic tools**

(A) ***neurog1**$^{hi1059}$*: Structure of the previously available *neurog1* allele from a retroviral insertion mutagenesis (Golling et al., 2002). The retroviral insertion leaves the open reading frame of Neurog1 intact. (B) ***neurog1**$^{m1469}$*: The *neurog1$^{m1469}$* allele was generated using CRISPR/Cas9 mediated genome editing. Two sgRNAs were designed and injected, marked by scissors, to induce a deletion of the whole bHLH domain coding sequence. The mutant allele can be identified by amplifying a 553 bp large fragment of the neurog1 ORF covering the bHLH domain. PCR on homozygous *neurog1$^{m1469}$* individuals results in a 396 bp shorter product. (C) ***olig2**$^{m1317}$*: The genomic transcribed region of the *olig2* locus is 2108 bp long and consists of two exons (grey) and one intron (dark yellow). The *olig2* open reading frame is located in exon 2 and is 822 bp long. The *olig2$^{m1317}$* mutant allele was generated using the TALEN mediated genome editing. The TALEN target site starts 324 bases after transcription start and is 45 bp long. It is located within the bHLH (blue) domain. The two TALEN sequences (cyan) flank the spacer sequence (yellow) which includes a HaeIII restriction site (red). The mutant allele has 13 bp deleted including the HaeIII restriction site and leads to a frameshift causing a preliminary stop codon (purple) at bp 355 – bp 357 in the bHLH domain and therefore to a truncated version of the protein. Genotyping is done by PCR followed by a HaeIII restriction digest. The forward primer (green) starts at the ATG of the coding sequence and the reverse primer (green) at bp 460 after transcription start. In wildtype, HaeIII digests the 460 bp fragment amplified by PCR into two fragments of 340 bp and 120 bp length. As the mutant is lacking the HaeIII restriction site, the amplified PCR fragment is not digested. Heterozygous fish have a mutant and a wildtype allele and therefore three PCR fragments are visible on the agarose gel after PCR and digest: the mutant uncut fragment of 460 bp and the two cut fragments of 340 bp and 120 bp length. (D) ***Tg(hsp70:olig2)m1306***: The open reading frame of *olig2* is put downstream of the hsp70l promoter and is followed by a SV40 polyadenylation site. For undemanding screening for the transgene, the construct contains GFP under the control of the *myl7* promoter. Abbreviations: AA, amino acid; D, digest; FP, forward primer; gDNA, genomic deoxyribonucleic acid; het, heterozygous; bHLH, basic helix-loop-helix domain; M, 100 bp marker; mut, mutant; ORF, open reading frame; polyA, SV40 polyadenylation sequence; RP, reverse primer; wt, wildtype.

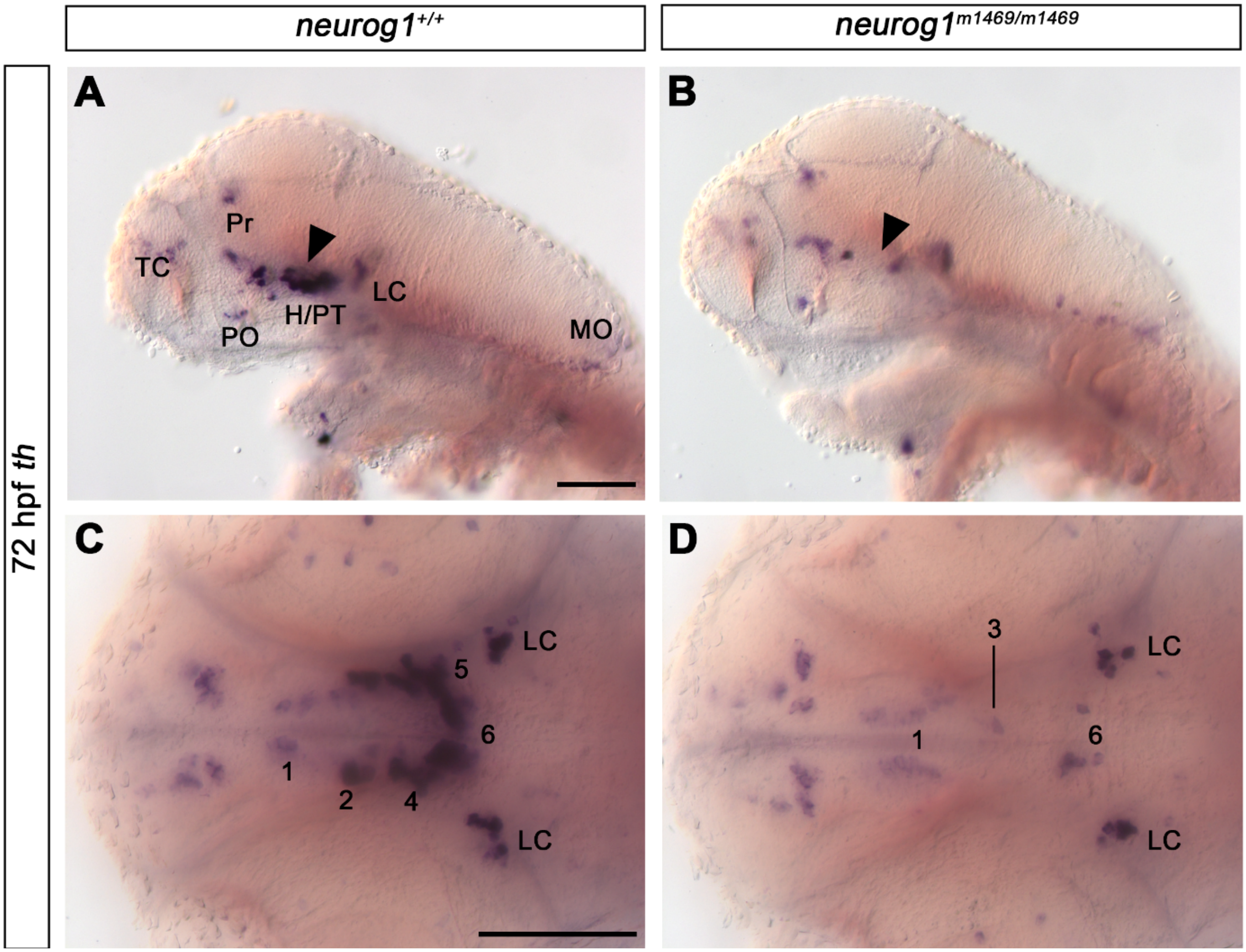


**Figure S2. *Th* expression in *neurog1*$^{m1469/m1469}$ mutants compared to wildtype siblings at 72 hpf.**
(A-D) *th* expression in (B,D) *neurog1*$^{m1469/m1469}$ mutants compared to (A,C) wildtype siblings at 72 hpf. In mutants DAC2, DAC4 and DAC5 (arrowheads in A and B) are missing and DAC6 is severely reduced. (E) (A,B) lateral views of enucleated embryos. (C,D) dorsal views. The phenotypes were observed in N/N (number of embryos with phenotype shown in figure panel / total number of embryos analyzed): (A,C) 7/7; (B,D) 9/9. Abbreviations: numbers 1 through 7 for DAC1-7, dopaminergic cell cluster 1-7; H, hypothalamus; LC, locus coeruleus; MO, medulla oblongata; PO, preoptic area; Pr, pretectum; PT, posterior tuberculum; TC, telencephalic cluster. Lateral views show single planes. Dorsal views show minimum projections of recorded stacks. Anterior is to the left. Scale bar: 100 µm.

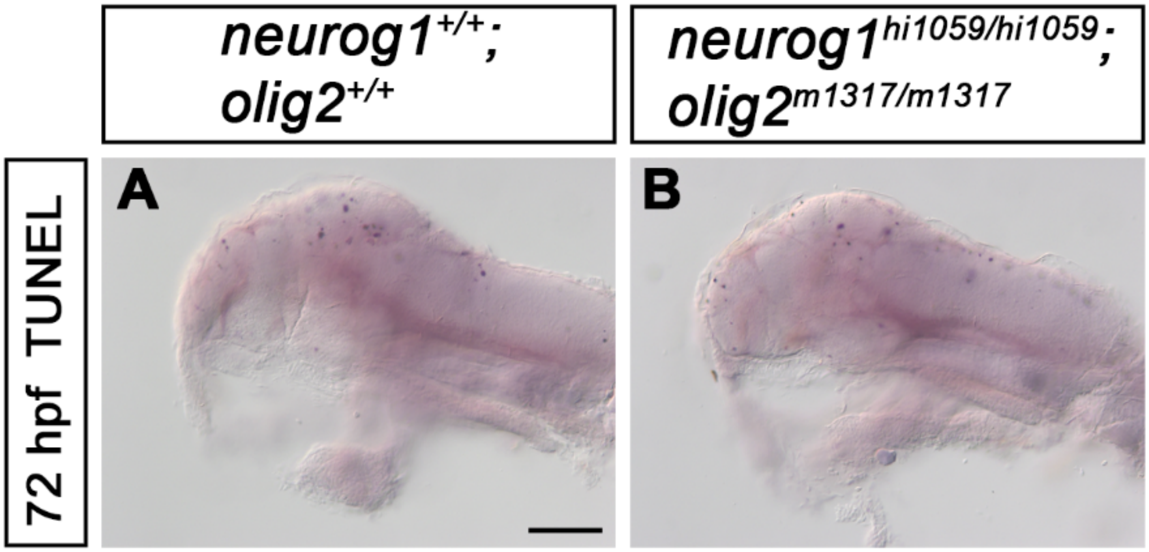


**Figure S3. *neurog1*$^{hi1059/hi1059}$ and *olig2*$^{m1317/m1317}$ double mutants do not show increased apoptosis.**
(A,B) TUNEL staining in (B) *neurog1*$^{hi1059/hi1059}$ and *olig2*$^{m1317/m1317}$ double mutants compared to (A) wildtype siblings. The phenotypes were observed in N/N (number of embryos with phenotype shown in figure panel / total number of embryos analyzed): (A) 5/5; (B) 10/10. Both images show single plane lateral views of enucleated embryos. Anterior is to the left. Scale bar: 100 µm.

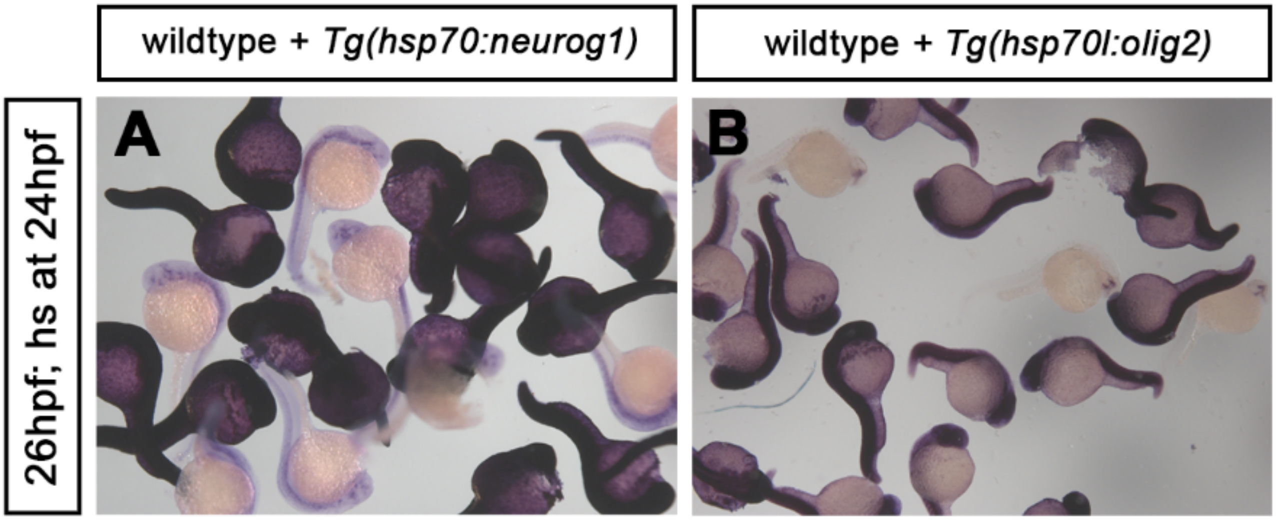


**Figure S4. Validation of *Tg(hsp70:neurog1)ups1* and *Tg(hsp70l:olig2)m1306* transgenic lines.**
An outcross of heterozygous *Tg(hsp70:neurog1)ups1* was heat-shocked, fixed soon after treatment and stained by WISH for *neurog1* mRNA, and an outcross of heterozygous *Tg(hsp70l:olig2)m1306* was heat-shocked, fixed soon after treatment and stained by WISH for *olig2* mRNA. Embryos displaying a strong, broad expression in the images are transgenic while embryos displaying the endogenous expression pattern are wild-type siblings.

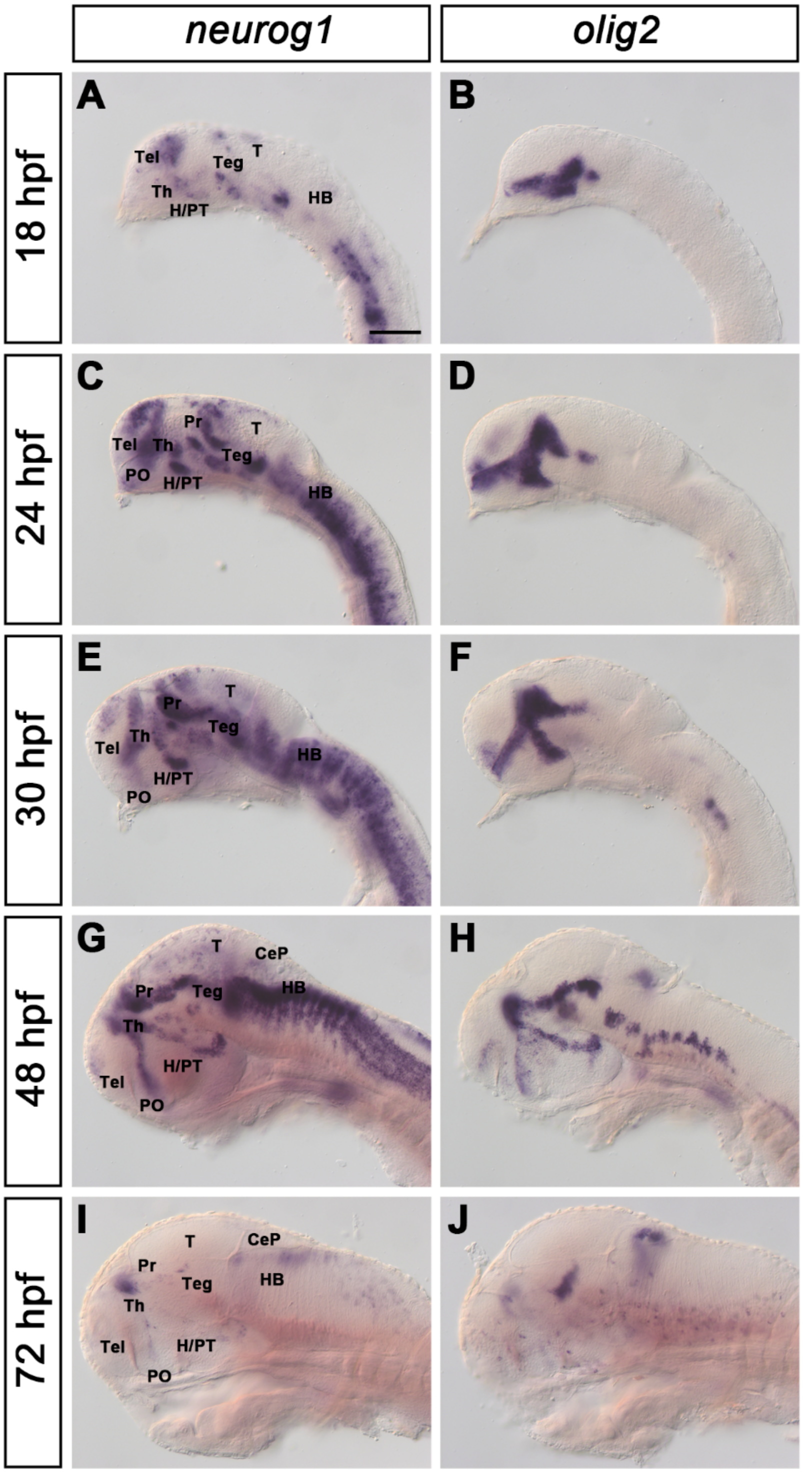


**Figure S5. Expression of *neurog1* and *olig2* during embryonic development.**

Lateral views of wildtype zebrafish larvae showing (A,C,E,G,I) *neurog1* and (B,D,F,H,J) *olig2* expression analyzed by whole mount *in situ* hybridization at (A,B) 18 hpf, (C,D) 24 hpf, (E,F) 30 hpf, (G,H) 48 hpf (*olig2*: enucleated) and (I,J) 72 hpf (enucleated embryos). Abbreviations: CeP, cerebellar plate; H, hypothalamus; HB, hindbrain; PO, preoptic area; Pr, pretectum; PT, posterior tuberculum; T, tectum; Teg, tegmentum; Tel, telencephalon; Th, thalamus. All images show single planes. Anterior is to the left. Scale bar: 100 µm.

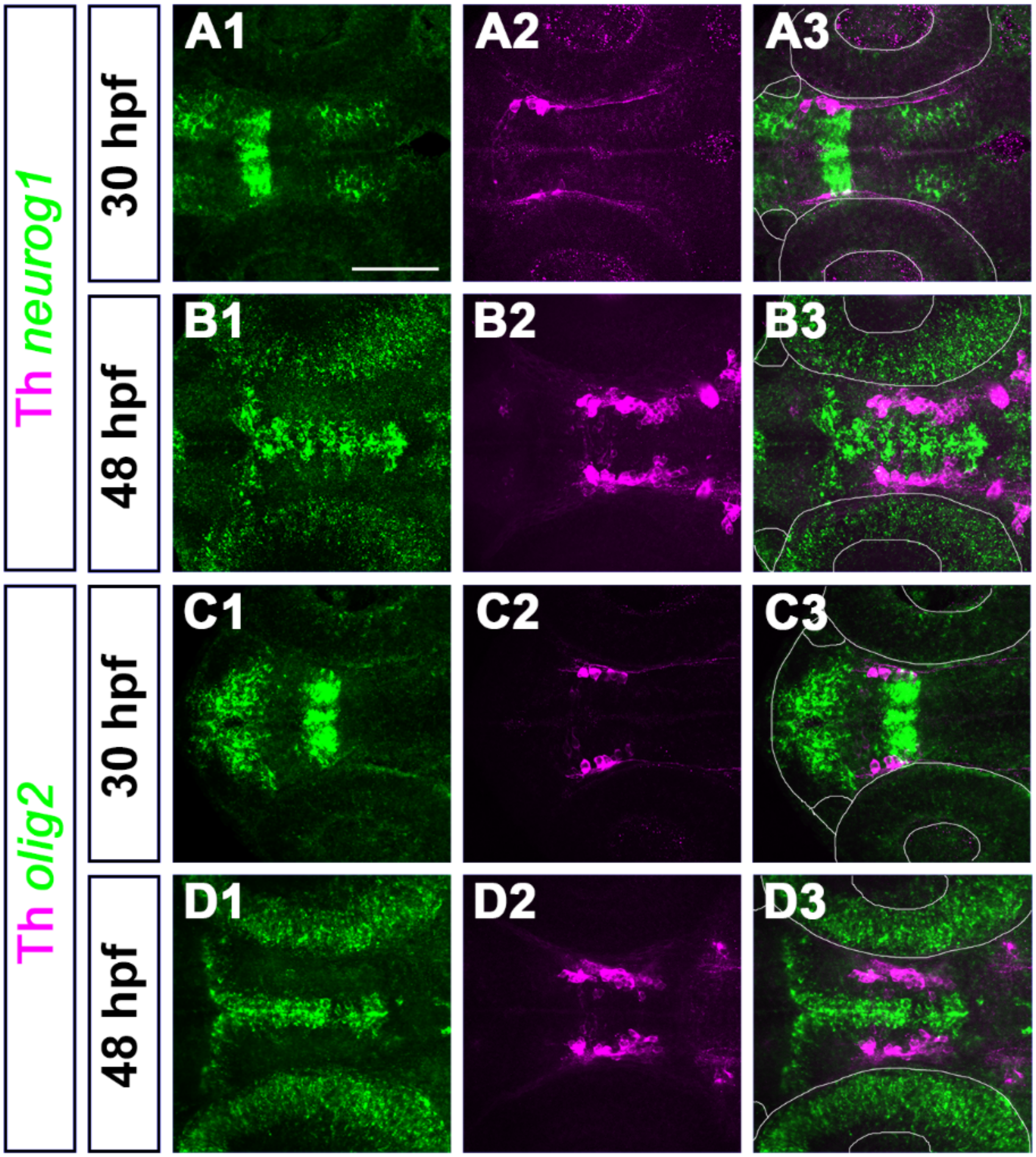


**Figure S6. *neurog1* and *olig2* are expressed in potential progenitors, but not in mature dopaminergic neurons.**

(A1-B3) Co-expression analysis of *neurog1* (green) and Th (magenta) at (A1-A3) 30 hpf and (B1-B3) 48 hpf. (C1-D3) co-expression of *olig2* (green) and Th (magenta) at (C1-C3) 30 hpf and (D1-D3) 48 hpf. At both stages, *neurog1* and *olig2* are not expressed in Th+ neurons. The white lines in (A3-D3) frame the outline of the imaged zebrafish larval head, nose pits and eyes; the line in (F2) frame the central nervous system of the zebrafish larval head. Dorsal views, anterior is to the left. All images are 15-35 µm maximum intensity projections. Scale bar: 100 µm.

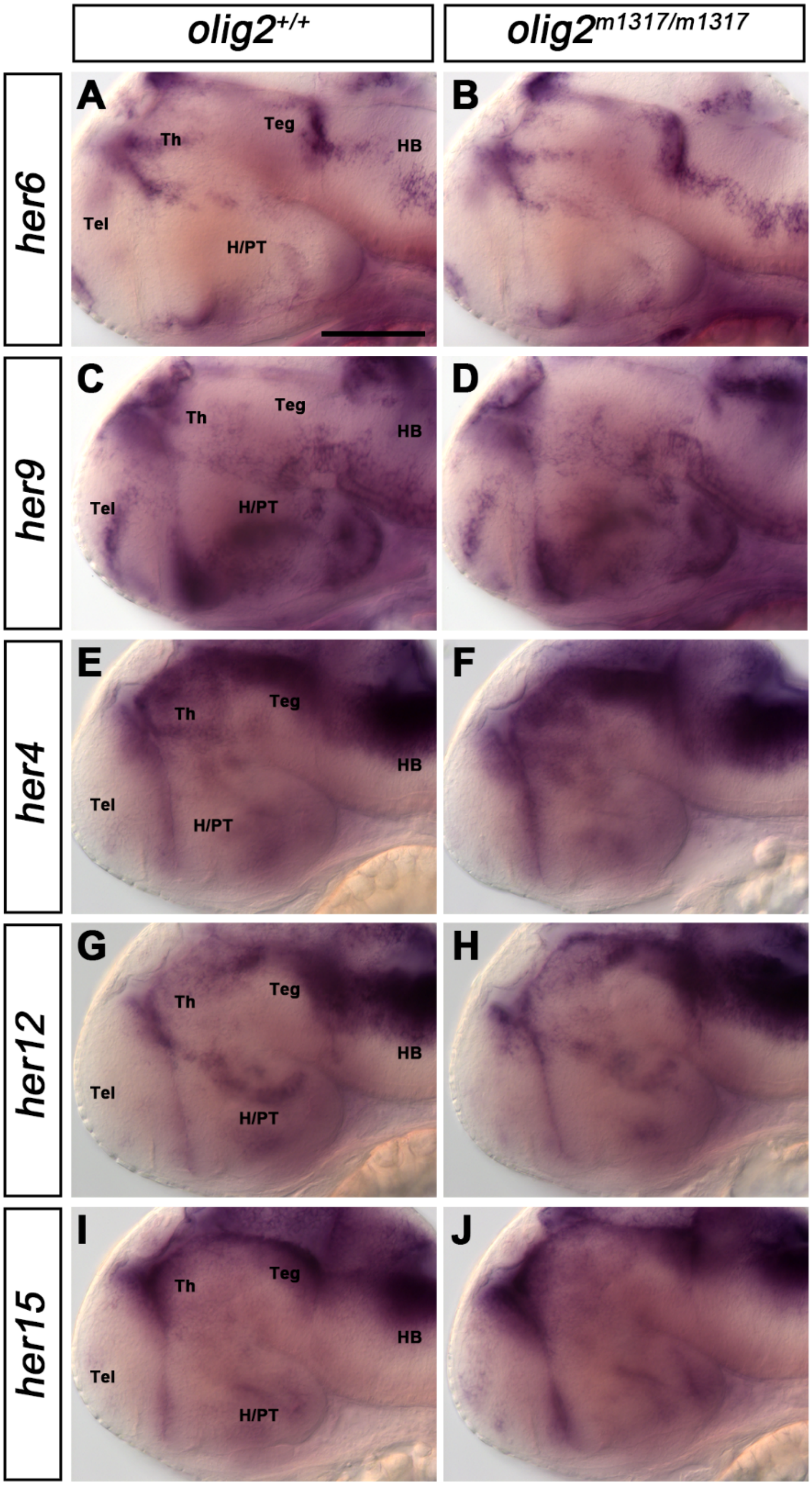


**Figure S7. Expression of *Hes1/Hes5* homologues in *olig2* mutants.**
(A,B) *her6* expression at 48 hpf in (B) *olig2*$^{m1317/m1317}$ mutant embryos compared to (A) wildtype siblings. (C,D) *her9* expression at 48 hpf in (D) *olig2*$^{m1317/m1317}$ mutant embryos compared to (C) wildtype siblings. (E,F) *her4* expression at 48 hpf in (F) *olig2*$^{m1317/m1317}$ mutant embryos compared to (E) wildtype siblings. (G,H) *her12* expression at 48 hpf in (H) *olig2*$^{m1317/m1317}$ mutant embryos compared to (G) wildtype siblings. (I,J) *her15* expression at 48 hpf in (J) *olig2*$^{m1317/m1317}$ mutant embryos compared to (I) wildtype siblings. The phenotypes were observed in N/N (number of embryos with phenotype shown in figure panel / total number of embryos analyzed): (A) 8/8; (B) 8/8; (C) 7/7; (D) 5/5; (E) 6/6; (F) 3/3; (G) 7/7; (H) 8/8; (I) 7/7; (J) 4/4. All images show lateral views and single planes. Anterior is to the left. Abbreviations: H, hypothalamus; HB - hindbrain; PT, posterior tuberculum; Teg, tegmentum; Tel, telencephalon; Th, thalamus. Scale bar: 100 µm.

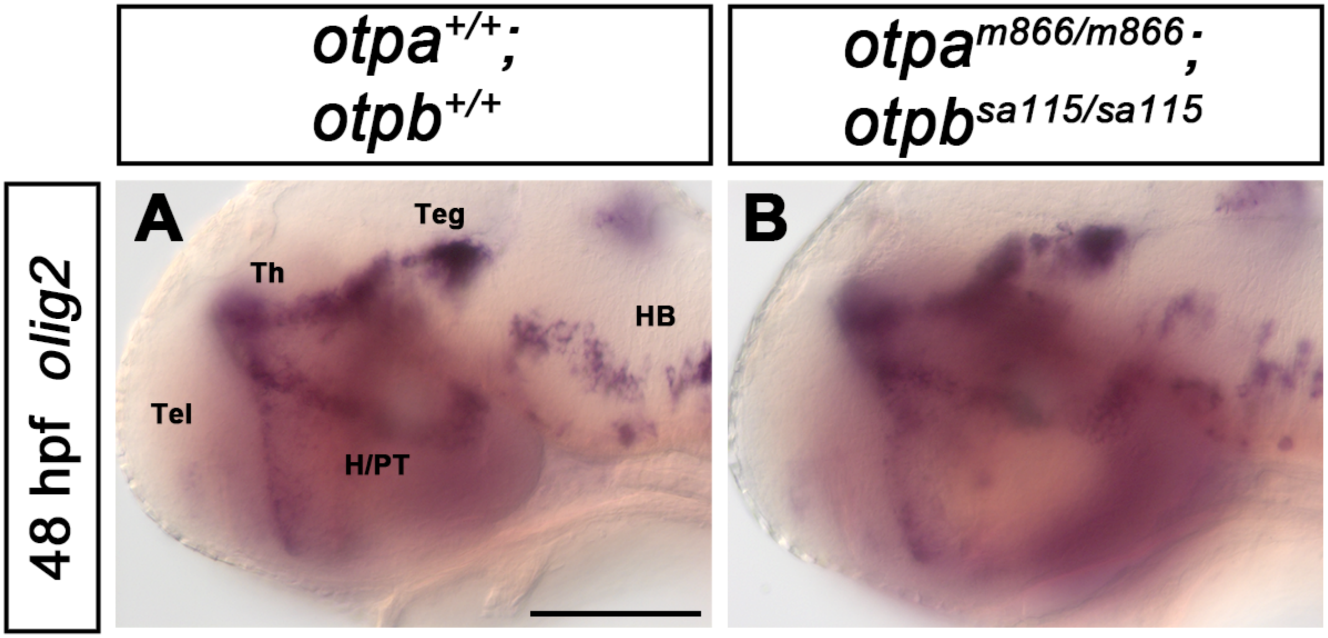


**Figure S8. Expression of *olig2* in *otpa*, *otpb* mutants**
(A,B) *olig2* expression at 48 hpf in (B) $otpa^{m866/m866}$; $otpb^{sa115/sa115}$ mutant embryos compared to (A) wildtype siblings. The phenotypes were observed in N/N (number of embryos with phenotype shown in figure panel / total number of embryos analyzed): (A) 4/4; (B) 5/5. All images show lateral views and single planes. Anterior is to the left. Levels were adjusted non-linearly, because embryos were not enucleated. Abbreviations: H, hypothalamus; HB - hindbrain; PT, posterior tuberculum; Teg, tegmentum; Tel, telencephalon; Th, thalamus. Scale bar: 100 µm.

#### Supplemental Movies

**Movie S1.** Dorsal view of *olig2* (green) and *neurog1* (magenta) co-expression analyzed by double fluorescent *in situ* hybridization at 30 hpf. Anterior is to the left. For anatomical orientation, see Figure 4. Figure 4A1-A3 is a maximum intensity projection of the planes 90-110. Scale bar: 50 µm.

**Movie S2.** Dorsal view of *olig2* (green) and *neurog1* (magenta) co-expression analyzed by double fluorescent *in situ* hybridization at 48 hpf. Anterior is to the left. For anatomical orientation, see Figure 4. Figure 4C1-C3 is a maximum intensity projection of the planes 95-134.Scale bar: 50 µm.

#### Supplemental Table 1

#### Cell Counts Figure 1

##### Figure 1 A,B

**30hpf neurog1 mut hi1059**

| embryo | DAC2&4 | genotype |
|---|---|---|
| 1 | 18 | wt |
| 2 | 20 | wt |
| 3 | 17 | het |
| 4 | 15 | het |
| 5 | 13 | het |
| 6 | 17 | het |
| 7 | 13 | het |
| 8 | 11 | het |
| 9 | 14 | het |
| 10 | 12 | het |
| embryo | DAC2&4 | genotype |
| 1 | 0 | neurog1 mut |
| 2 | 0 | neurog1 mut |
| 3 | 2 | neurog1 mut |
| 4 | 2 | neurog1 mut |
| 5 | 1 | neurog1 mut |
| 6 | 2 | neurog1 mut |
| 7 | 3 | neurog1 mut |
| 8 | 0 | neurog1 mut |

| | Total |
|---|---|
| Mean WT/HET | 15,000 |
| SE WT/HET | 0,919 |
| Mean MUT | 1,250 |
| SE MUT | 0,412 |
| P Value | 0,0002 |

### Figure 1 M

**72hpf neurog1 mut hi1059**

| embryo | DAC1 | DAC2 | DAC3 | DAC4 | DAC5 | DAC6 | LC | PO | Pr | SP | OB | genotype |
|---|---|---|---|---|---|---|---|---|---|---|---|---|
| 1 | 35 | 12 | 7 | 10 | 41 | 31 | 16 | 1 | 12 | 3 | 0 | wt |
| 2 | 41 | 10 | 3 | 13 | 34 | 44 | 11 | 5 | 22 | 3 | 0 | wt |
| 3 | 28 | 12 | 6 | 12 | 34 | 35 | 11 | 3 | 13 | 3 | 0 | wt |
| 4 | 34 | 13 | 8 | 14 | 40 | 42 | 11 | 10 | 15 | 9 | 3 | wt |
| 5 | 31 | 11 | 8 | 12 | 39 | 43 | 12 | 9 | 25 | 5 | 3 | wt |
| 6 | 19 | 10 | 7 | 11 | 42 | 42 | 14 | 6 | 29 | 10 | 5 | wt |
| 7 | 32 | 10 | 10 | 14 | 40 | 50 | 11 | 4 | 22 | 8 | 5 | wt |
| 8 | 43 | 12 | 7 | 11 | 34 | 38 | 14 | 5 | 31 | 8 | 6 | wt |
| embryo | DAC1 | DAC2 | DAC3 | DAC4 | DAC5 | DAC6 | LC | PO | Pr | SP | OB | genotype |
| 1 | 23 | 0 | 7 | 0 | 0 | 25 | 14 | 11 | 12 | 5 | 0 | neurog1 mut |
| 2 | 32 | 0 | 6 | 0 | 0 | 28 | 11 | 5 | 24 | 3 | 5 | neurog1 mut |
| 3 | 31 | 0 | 11 | 0 | 0 | 25 | 15 | 5 | 23 | 8 | 4 | neurog1 mut |
| 4 | 28 | 0 | 6 | 0 | 0 | 21 | 14 | 8 | 28 | 10 | 2 | neurog1 mut |
| 5 | 31 | 0 | 10 | 0 | 0 | 19 | 14 | 3 | 28 | 6 | 4 | neurog1 mut |
| 6 | 34 | 0 | 5 | 0 | 0 | 27 | 15 | 5 | 14 | 7 | 2 | neurog1 mut |
| 7 | 37 | 0 | 9 | 0 | 0 | 20 | 15 | 11 | 22 | 15 | 10 | neurog1 mut |
| 8 | 34 | 0 | 5 | 0 | 0 | 26 | 14 | 7 | 21 | 10 | 6 | neurog1 mut |
| 9 | 28 | 1 | 7 | 0 | 0 | 21 | 14 | 10 | 16 | 7 | 3 | neurog1 mut |
| 10 | 25 | 0 | 7 | 0 | 0 | 20 | 13 | 9 | 20 | 5 | 3 | neurog1 mut |
| 11 | 32 | 0 | 14 | 0 | 0 | 19 | 14 | 5 | 20 | 10 | 3 | neurog1 mut |
| 12 | 27 | 0 | 12 | 0 | 0 | 19 | 13 | 9 | 16 | 7 | 1 | neurog1 mut |

| | DAC2 | DAC4 | DAC5 | DAC6 | DAC1 | DAC3 | LC | PO | Pr | SP | OB |
|---|---|---|---|---|---|---|---|---|---|---|---|
| Mean WT | 11,250 | 12,125 | 38,000 | 40,625 | 32,875 | 7,000 | 12,500 | 5,375 | 21,125 | 6,125 | 2,750 |
| SE WT | 0,412 | 0,515 | 1,210 | 2,070 | 2,655 | 0,707 | 0,681 | 1,051 | 2,546 | 1,043 | 0,881 |
| Mean MUT | 0,083 | 0,000 | 0,000 | 22,500 | 30,167 | 8,250 | 13,833 | 7,333 | 20,333 | 7,750 | 3,583 |
| SE MUT | 0,083 | 0,000 | 0,000 | 0,989 | 1,173 | 0,845 | 0,322 | 0,782 | 1,479 | 0,914 | 0,753 |
| P Values | < 0.0001 | < 0.0001 | < 0.0001 | < 0.0001 | 0,2671 | 0,6622 | 0,0934 | 0,1686 | 0,7482 | 0,3675 | 0,6512 |

### Figure 1 G,H

**30hpf** **olig2 mut** **m1317**

| embryo | total | genotype |
|---|---|---|
| 1 | 30 | wt |
| 2 | 22 | wt |
| 3 | 25 | wt |
| 4 | 25 | wt |
| 5 | 21 | wt |
| 6 | 28 | wt |
| embryo | total | genotype |
| 1 | 21 | olig2 mut |
| 2 | 15 | olig2 mut |
| 3 | 10 | olig2 mut |
| 4 | 18 | olig2 mut |
| 5 | 14 | olig2 mut |
| 6 | 21 | olig2 mut |

| | total |
|---|---|
| Mean WT/HET | 25,167 |
| SE WT/HET | 1,400 |
| Mean MUT | 16,500 |
| SE MUT | 1,765 |
| P Value | 0,0065 |

### Figure 1 N

**72hpf olig2 mut m1317**

| embryo | DAC1 | DAC2 | DAC3 | DAC4 | DAC5 | DAC6 | LC | PO | Pr | SP | OB | genotype |
|---|---|---|---|---|---|---|---|---|---|---|---|---|
| 1 | 52 | 9 | 10 | 15 | 41 | 29 | 12 | 18 | 29 | 9 | 20 | wt |
| 2 | 38 | 11 | 11 | 15 | 36 | 47 | 10 | 17 | 31 | 17 | 14 | wt |
| 3 | 63 | 12 | 9 | 12 | 37 | 35 | 12 | 15 | 46 | 12 | 12 | wt |
| 4 | 39 | 13 | 8 | 15 | 40 | 37 | 15 | 13 | 32 | 13 | 12 | wt |
| 5 | 58 | 11 | 14 | 19 | 37 | 40 | 11 | 15 | 45 | 10 | 16 | wt |
| 6 | 54 | 16 | 8 | 19 | 40 | 41 | 14 | 13 | 45 | 9 | 11 | wt |
| 7 | 48 | 10 | 15 | 19 | 41 | 42 | 10 | 17 | 39 | 11 | 10 | wt |
| embryo | DAC1 | DAC2 | DAC3 | DAC4 | DAC5 | DAC6 | LC | PO | Pr | SP | OB | genotype |
| 1 | 43 | 6 | 14 | 18 | 30 | 8 | 11 | 9 | 21 | 13 | 12 | olig2 mut |
| 2 | 50 | 6 | 18 | 16 | 37 | 3 | 16 | 19 | 41 | 9 | 16 | olig2 mut |
| 3 | 58 | 5 | 12 | 14 | 48 | 3 | 12 | 8 | 39 | 5 | 9 | olig2 mut |
| 4 | 39 | 10 | 11 | 11 | 28 | 9 | 15 | 14 | 32 | 4 | 9 | olig2 mut |
| 5 | 36 | 4 | 21 | 19 | 41 | 1 | 20 | 12 | 32 | 13 | 13 | olig2 mut |
| 6 | 42 | 4 | 18 | 17 | 36 | 3 | 20 | 15 | 45 | 16 | 16 | olig2 mut |
| 7 | 43 | 4 | 7 | 14 | 28 | 6 | 14 | 10 | 24 | 14 | 18 | olig2 mut |

| | DAC2 | DAC4 | DAC5 | DAC6 | DAC1 | DAC3 | LC | PO | Pr | SP | OB |
|---|---|---|---|---|---|---|---|---|---|---|---|
| Mean WT | 11,714 | 16,286 | 38,857 | 38,714 | 50,286 | 10,714 | 12,000 | 15,429 | 38,143 | 11,571 | 13,571 |
| SE WT | 0,865 | 1,040 | 0,800 | 2,168 | 3,523 | 1,063 | 0,724 | 0,751 | 2,798 | 1,066 | 1,307 |
| Mean MUT | 5,571 | 15,571 | 35,429 | 4,714 | 44,429 | 14,429 | 15,429 | 12,429 | 33,429 | 10,571 | 13,286 |
| SE MUT | 0,812 | 1,043 | 2,810 | 1,128 | 2,785 | 1,837 | 1,343 | 1,462 | 3,344 | 1,757 | 1,340 |
| P Values | 0,0017 | 0,542 | 0,2611 | 0,0006 | 0,2972 | 0,1649 | 0,0594 | 0,1206 | 0,3986 | > 0.9999 | 0,9534 |

#### Supplemental Table 2

#### Cell Counts Figure 2 I

**Control treatment** **hi1059+m1317**

| embryo | DAC1 | DAC2 | DAC3 | DAC4 | DAC5 | DAC6 | LC | PO | genotype |
|---|---|---|---|---|---|---|---|---|---|
| 1 | 20 | 12 | 8 | 12 | 33 | 64 | 14 | 5 | wt |
| 2 | 11 | 13 | 6 | 11 | 23 | 50 | 9 | 6 | wt |
| 3 | 9 | 10 | 10 | 13 | 37 | 54 | 8 | 6 | wt |
| 4 | 17 | 8 | 5 | 12 | 35 | 56 | 11 | 11 | wt |
| 5 | 17 | 9 | 8 | 9 | 36 | 60 | 12 | 7 | wt |
| 6 | 7 | 10 | 7 | 7 | 40 | 57 | 11 | 5 | wt |
| 7 | 11 | 11 | 7 | 9 | 35 | 57 | 12 | 4 | wt |
| embryo | DAC1 | DAC2 | DAC3 | DAC4 | DAC5 | DAC6 | LC | PO | genotype |
| 1 | 11 | 0 | 10 | 0 | 0 | 24 | 11 | 5 | neurog1 mut |
| 2 | 20 | 0 | 7 | 0 | 0 | 29 | 8 | 4 | neurog1 mut |
| 3 | 16 | 0 | 5 | 0 | 0 | 23 | 11 | 2 | neurog1 mut |
| 4 | 11 | 0 | 6 | 0 | 0 | 22 | 11 | 6 | neurog1 mut |
| 5 | 8 | 0 | 10 | 0 | 0 | 23 | 15 | 5 | neurog1 mut |
| 6 | 15 | 0 | 9 | 0 | 0 | 29 | 9 | 13 | neurog1 mut |
| embryo | DAC1 | DAC2 | DAC3 | DAC4 | DAC5 | DAC6 | LC | PO | genotype |
| 1 | 14 | 4 | 9 | 9 | 35 | 12 | 19 | 6 | olig2 mut |
| 2 | 17 | 3 | 6 | 13 | 29 | 19 | 9 | 7 | olig2 mut |
| 3 | 15 | 4 | 5 | 10 | 30 | 23 | 8 | 5 | olig2 mut |
| 4 | 12 | 3 | 7 | 11 | 38 | 17 | 10 | 10 | olig2 mut |
| 5 | 9 | 5 | 6 | 9 | 27 | 22 | 8 | 4 | olig2 mut |
| 6 | 8 | 3 | 8 | 8 | 29 | 15 | 12 | 5 | olig2 mut |
| 7 | 7 | 2 | 12 | 11 | 30 | 23 | 12 | 6 | olig2 mut |
| embryo | DAC1 | DAC2 | DAC3 | DAC4 | DAC5 | DAC6 | LC | PO | genotype |
| 1 | 7 | 0 | 3 | 0 | 0 | 12 | 12 | 7 | double mut |
| 2 | 17 | 0 | 8 | 0 | 0 | 7 | 9 | 8 | double mut |
| 3 | 10 | 0 | 10 | 0 | 0 | 16 | 11 | 5 | double mut |
| 4 | 15 | 0 | 10 | 0 | 0 | 10 | 15 | 8 | double mut |
| 5 | 13 | 0 | 12 | 0 | 0 | 11 | 13 | 10 | double mut |
| 6 | 9 | 0 | 5 | 0 | 0 | 10 | 11 | 6 | double mut |
| 7 | 9 | 0 | 5 | 0 | 0 | 12 | 12 | 6 | double mut |
| 8 | 12 | 0 | 8 | 0 | 0 | 14 | 14 | 8 | double mut |
| 9 | 12 | 0 | 5 | 0 | 0 | 14 | 10 | 6 | double mut |
| 10 | 15 | 0 | 10 | 0 | 0 | 5 | 10 | 5 | double mut |
| 11 | 11 | 0 | 3 | 0 | 0 | 14 | 12 | 4 | double mut |
| 12 | 15 | 0 | 13 | 0 | 0 | 9 | 8 | 8 | double mut |

**DAPT treatment hi1059+m1317**

| embryo | DAC1 | DAC2 | DAC3 | DAC4 | DAC5 | DAC6 | LC | PO | genotype |
|---|---|---|---|---|---|---|---|---|---|
| 1 | 15 | 10 | 0 | 16 | 60 | 74 | 11 | 2 | wt |
| 2 | 8 | 7 | 0 | 10 | 66 | 70 | 12 | 7 | wt |
| 3 | 3 | 7 | 0 | 9 | 59 | 80 | 15 | 6 | wt |
| 5 | 5 | 9 | 0 | 11 | 54 | 66 | 10 | 4 | wt |
| 6 | 9 | 6 | 3 | 11 | 52 | 74 | 10 | 1 | wt |
| 7 | 10 | 10 | 1 | 10 | 70 | 68 | 14 | 5 | wt |
| 8 | 19 | 11 | 1 | 10 | 47 | 58 | 12 | 4 | wt |
| 9 | 6 | 10 | 0 | 12 | 49 | 53 | 9 | 6 | wt |
| 10 | 4 | 16 | 3 | 15 | 62 | 60 | 10 | 6 | wt |
| 11 | 9 | 7 | 5 | 8 | 55 | 62 | 13 | 5 | wt |
| embryo | DAC1 | DAC2 | DAC3 | DAC4 | DAC5 | DAC6 | LC | PO | genotype |
| 1 | 13 | 0 | 2 | 0 | 0 | 27 | 8 | 3 | neurog1 mut |
| 2 | 5 | 0 | 2 | 0 | 0 | 22 | 13 | 5 | neurog1 mut |
| 3 | 8 | 0 | 0 | 0 | 0 | 27 | 9 | 7 | neurog1 mut |
| 4 | 9 | 0 | 3 | 0 | 0 | 24 | 14 | 5 | neurog1 mut |
| 5 | 7 | 0 | 2 | 0 | 0 | 27 | 12 | 8 | neurog1 mut |
| 6 | 12 | 0 | 2 | 0 | 0 | 20 | 11 | 4 | neurog1 mut |
| 7 | 10 | 0 | 1 | 0 | 0 | 25 | 14 | 5 | neurog1 mut |
| 8 | 7 | 0 | 3 | 0 | 0 | 24 | 15 | 5 | neurog1 mut |
| embryo | DAC1 | DAC2 | DAC3 | DAC4 | DAC5 | DAC6 | LC | PO | genotype |
| 1 | 5 | 1 | 3 | 8 | 39 | 42 | 11 | 3 | olig2 mut |
| 2 | 14 | 0 | 0 | 13 | 33 | 24 | 15 | 4 | olig2 mut |
| 3 | 5 | 1 | 1 | 7 | 24 | 47 | 13 | 5 | olig2 mut |
| 4 | 6 | 0 | 1 | 11 | 38 | 36 | 11 | 5 | olig2 mut |
| 5 | 13 | 0 | 2 | 10 | 32 | 35 | 13 | 6 | olig2 mut |
| 6 | 4 | 0 | 3 | 8 | 31 | 45 | 8 | 7 | olig2 mut |
| 7 | 8 | 0 | 2 | 12 | 30 | 54 | 13 | 7 | olig2 mut |
| embryo | DAC1 | DAC2 | DAC3 | DAC4 | DAC5 | DAC6 | LC | PO | genotype |
| 1 | 2 | 0 | 0 | 0 | 0 | 11 | 12 | 4 | double mut |
| 2 | 2 | 0 | 0 | 0 | 0 | 10 | 14 | 3 | double mut |
| 3 | 17 | 0 | 4 | 0 | 0 | 16 | 15 | 6 | double mut |
| 4 | 8 | 0 | 2 | 0 | 0 | 11 | 14 | 7 | double mut |
| 5 | 6 | 0 | 3 | 0 | 0 | 5 | 16 | 10 | double mut |
| 6 | 5 | 0 | 0 | 0 | 0 | 10 | 12 | 4 | double mut |
| 7 | 6 | 0 | 0 | 0 | 0 | 16 | 12 | 5 | double mut |
| 8 | 6 | 0 | 0 | 0 | 0 | 14 | 11 | 8 | double mut |
| 9 | 11 | 0 | 4 | 0 | 0 | 9 | 12 | 6 | double mut |
| 10 | 11 | 0 | 1 | 0 | 0 | 5 | 10 | 3 | double mut |

| MEAN | DAC1 | DAC2 | DAC3 | DAC4 | DAC5 | DAC6 | LC | PO |
|---|---|---|---|---|---|---|---|---|
| wt CTR | 13,14 | 10,43 | 7,29 | 10,43 | 34,14 | 56,86 | 11,00 | 6,29 |
| neurog1 mut CTR | 13,50 | 0,00 | 7,83 | 0,00 | 0,00 | 25,00 | 10,83 | 5,83 |
| olig2 mut CTR | 11,71 | 3,43 | 7,57 | 10,14 | 31,14 | 18,71 | 11,14 | 6,14 |
| double mut CTR | 12,08 | 0,00 | 7,67 | 0,00 | 0,00 | 11,17 | 11,42 | 6,75 |
| wt DAPT | 8,80 | 9,30 | 1,30 | 11,20 | 57,40 | 66,50 | 11,60 | 4,60 |
| neurog1 mut DAPT | 8,88 | 0,00 | 1,88 | 0,00 | 0,00 | 24,50 | 12,00 | 5,25 |
| olig2 mut DAPT | 7,86 | 0,29 | 1,71 | 9,86 | 32,43 | 40,43 | 12,00 | 5,29 |
| double mut DAPT | 7,40 | 0,00 | 1,40 | 0,00 | 0,00 | 10,70 | 12,80 | 5,60 |

| SE | DAC1 | DAC2 | DAC3 | DAC4 | DAC5 | DAC6 | LC | PO |
|---|---|---|---|---|---|---|---|---|
| wt CTR | 1,831 | 0,649 | 0,606 | 0,812 | 2,029 | 1,668 | 0,756 | 0,865 |
| neurog1 mut CTR | 1,765 | 0,000 | 0,872 | 0,000 | 0,000 | 1,291 | 0,980 | 1,537 |
| olig2 mut CTR | 1,443 | 0,369 | 0,896 | 0,634 | 1,471 | 1,614 | 1,455 | 0,738 |
| double mut CTR | 0,874 | 0,000 | 0,987 | 0,000 | 0,000 | 0,920 | 0,583 | 0,494 |
| wt DAPT | 1,576 | 0,920 | 0,559 | 0,800 | 2,330 | 2,638 | 0,618 | 0,600 |
| neurog1 mut DAPT | 0,953 | 0,000 | 0,350 | 0,000 | 0,000 | 0,906 | 0,886 | 0,559 |
| olig2 mut DAPT | 0,953 | 0,000 | 0,350 | 0,000 | 0,000 | 0,906 | 0,886 | 0,559 |
| double mut DAPT | 1,447 | 0,000 | 0,542 | 0,000 | 0,000 | 1,230 | 0,593 | 0,718 |

| P Values - CTR | DAC2 | DAC4 | DAC5 | DAC6 |
|---|---|---|---|---|
| wt vs neurog1 mut | 0,0012 | 0,0012 | 0,0012 | 0,0012 |
| wt vs olig2 mut | 0,0006 | 0,7372 | 0,197 | 0,0006 |
| wt vs double mut | <0,0001 | <0,0001 | <0,0001 | <0,0001 |
| neurog1 mut vs olig2 mut | 0,0012 | 0,0012 | 0,0012 | 0,0175 |
| neurog1 mut vs double mut | 0,9999 | 0,9999 | 0,9999 | 0,0001 |
| olig2 mut vs double mut | <0,0001 | <0,0001 | <0,0001 | 0,0011 |

| P Values - DAPT | DAC2 | DAC4 | DAC5 | DAC6 |
|---|---|---|---|---|
| wt vs neurog1 mut | <0,0001 | <0,0001 | <0,0001 | <0,0001 |
| wt vs olig2 mut | <0,0001 | 0,3972 | 0,0001 | 0,0002 |
| wt vs double mut | <0,0001 | <0,0001 | <0,0001 | <0,0001 |
| neurog1 mut vs olig2 mut | 0,2 | 0,0002 | 0,0002 | 0,0062 |
| neurog1 mut vs double mut | 0,9999 | 0,9999 | 0,9999 | <0,0001 |
| olig2 mut vs double mut | 0,1544 | <0,0001 | <0,0001 | 0,0001 |

| P Values - CTR vs DAPT | DAC2 | DAC4 | DAC5 | DAC6 |
|---|---|---|---|---|
| wt | 0,2088 | 0,8073 | 0,0001 | 0,0143 |
| neurog1 mut | 0,9999 | 0,9999 | 0,9999 | 0,976 |
| olig2 mut | 0,0006 | 0,7762 | 0,3304 | 0,0006 |
| double mut | 0,9999 | 0,9999 | 0,9999 | 0,7583 |

#### Supplemental Table 3

#### Embryos Figure 3 A,B

| hsp70:neurog1 | ups1 | | | |
|---|---|---|---|---|
| genotype | treatment | stage (hpf) | phenotype | embryos evaluated |
| wt | hs at 12hpf | 30hpf | similar to untreated | (5/6) |
| wt | hs at 12hpf | 72hpf | similar to untreated | (6/6) |
| hsp70:neurog1 | hs at 12hpf | 30hpf | increased stain intensity | (6/6) |
| hsp70:neurog1 | hs at 12hpf | 72hpf | increased stain intensity | (6/6) |

#### Cell Counts Figure 3 G,H

| 30 hpf | hsp70:olig2 | m1306 |
|---|---|---|
| embryo | total | genotype |
| 1 | 20 | wt |
| 2 | 15 | wt |
| 3 | 19 | wt |
| 4 | 19 | wt |
| 5 | 23 | wt |
| 6 | 26 | wt |
| 7 | 19 | wt |
| 8 | 21 | wt |
| 9 | 19 | wt |
| 10 | 17 | wt |
| 11 | 26 | wt |
| 12 | 9 | wt |
| embryo | total | genotype |
| 1 | 31 | hsp70:olig2 |
| 2 | 32 | hsp70:olig2 |
| 3 | 29 | hsp70:olig2 |
| 4 | 31 | hsp70:olig2 |
| 5 | 20 | hsp70:olig2 |
| 6 | 31 | hsp70:olig2 |
| 7 | 25 | hsp70:olig2 |
| 8 | 28 | hsp70:olig2 |
| 9 | 24 | hsp70:olig2 |
| 10 | 32 | hsp70:olig2 |
| 11 | 40 | hsp70:olig2 |
| 12 | 38 | hsp70:olig2 |

| | total |
|---|---|
| Mean WT | 19,417 |
| SE WT | 1,340 |
| Mean TG | 30,083 |
| SE TG | 1,607 |
| P Value | < 0.0001 |

### Cell counts Figure 3 M

**72 hpf hsp70:olig2 m1306**

| embryo | DAC1 | DAC2 | DAC3 | DAC4 | DAC5 | DAC6 | LC | PO | Pr | SP | OB | genotype |
|---|---|---|---|---|---|---|---|---|---|---|---|---|
| 1 | 46 | 11 | 19 | 20 | 29 | 43 | 15 | 9 | 31 | 13 | 13 | wt |
| 2 | 40 | 9 | 8 | 16 | 31 | 31 | 14 | 11 | 25 | 17 | 27 | wt |
| 3 | 24 | 13 | 6 | 10 | 35 | 36 | 15 | 10 | 11 | 10 | 9 | wt |
| 4 | 44 | 9 | 13 | 13 | 44 | 35 | 14 | 8 | 29 | 14 | 11 | wt |
| 5 | 49 | 11 | 8 | 12 | 34 | 37 | 16 | 13 | 35 | 12 | 13 | wt |
| 6 | 58 | 10 | 13 | 14 | 32 | 47 | 11 | 15 | 40 | 14 | 25 | wt |
| 7 | 43 | 13 | 21 | 14 | 26 | 37 | 13 | 14 | 30 | 17 | 16 | wt |
| 8 | 38 | 10 | 12 | 16 | 29 | 37 | 17 | 14 | 29 | 9 | 16 | wt |
| 9 | 58 | 12 | 12 | 14 | 34 | 39 | 14 | 10 | 35 | 11 | 17 | wt |
| 10 | 37 | 10 | 20 | 16 | 29 | 37 | 12 | 8 | 21 | 14 | 21 | wt |
| 11 | 48 | 10 | 9 | 13 | 35 | 47 | 14 | 12 | 18 | 11 | 18 | wt |
| 12 | 34 | 14 | 16 | 13 | 46 | 47 | 16 | 19 | 35 | 19 | 19 | wt |
| embryo | DAC1 | DAC2 | DAC3 | DAC4 | DAC5 | DAC6 | LC | PO | Pr | SP | OB | genotype |
| 1 | 45 | 19 | 11 | 10 | 35 | 34 | 0 | 6 | 45 | 17 | 23 | hs:olig2 |
| 2 | 41 | 18 | 6 | 10 | 29 | 44 | 0 | 6 | 15 | 11 | 8 | hs:olig2 |
| 3 | 32 | 20 | 10 | 10 | 36 | 38 | 0 | 9 | 23 | 17 | 17 | hs:olig2 |
| 4 | 38 | 19 | 9 | 10 | 35 | 40 | 0 | 13 | 29 | 8 | 8 | hs:olig2 |
| 5 | 43 | 25 | 8 | 12 | 29 | 36 | 0 | 9 | 20 | 13 | 17 | hs:olig2 |
| 6 | 41 | 25 | 13 | 11 | 30 | 39 | 0 | 11 | 22 | 8 | 15 | hs:olig2 |
| 7 | 43 | 18 | 19 | 15 | 32 | 40 | 0 | 14 | 28 | 10 | 11 | hs:olig2 |
| 8 | 46 | 17 | 10 | 13 | 38 | 48 | 0 | 10 | 26 | 16 | 9 | hs:olig2 |
| 9 | 48 | 24 | 8 | 15 | 30 | 52 | 0 | 7 | 26 | 11 | 5 | hs:olig2 |
| 10 | 37 | 20 | 13 | 13 | 29 | 48 | 0 | 12 | 44 | 20 | 18 | hs:olig2 |
| 11 | 48 | 23 | 9 | 15 | 34 | 47 | 0 | 11 | 34 | 15 | 17 | hs:olig2 |
| 12 | 35 | 18 | 13 | 12 | 34 | 47 | 0 | 8 | 41 | 13 | 17 | hs:olig2 |

| | DAC1 | DAC2 | DAC3 | DAC4 | DAC5 | DAC6 | LC | PO | Pr | SP | OB |
|---|---|---|---|---|---|---|---|---|---|---|---|
| Mean WT | 43,250 | 11,000 | 13,083 | 14,250 | 33,667 | 39,417 | 14,250 | 11,917 | 28,250 | 13,417 | 17,083 |
| SE WT | 2,796 | 0,477 | 1,443 | 0,730 | 1,734 | 1,535 | 0,494 | 0,941 | 2,387 | 0,883 | 1,554 |
| Mean TG | 41,417 | 20,500 | 10,750 | 12,167 | 32,583 | 42,750 | 0,000 | 9,667 | 29,417 | 13,250 | 13,750 |
| SE TG | 1,474 | 0,848 | 0,986 | 0,588 | 0,908 | 1,634 | 0,000 | 0,762 | 2,792 | 1,102 | 1,558 |
| P Values | 0,5597 | < 0.0001 | 0,3089 | 0,1169 | 0,9656 | 0,1071 | < 0.0001 | 0,1082 | n.d. | 0,8532 | 0,2002 |
